# Bat teeth illuminate the diversification of mammalian tooth classes

**DOI:** 10.1101/2021.12.05.471324

**Authors:** Alexa Sadier, Neal Anthwal, Andrew L. Krause, Renaud Dessalles, Michael Lake, Laurent Bentolila, Robert Haase, Natalie Nieves, Sharlene Santana, Karen Sears

## Abstract

Tooth classes are a mammalian innovation that has contributed to the evolutionary success of mammals. However, our understanding of how tooth classes have evolved and diversified remains limited. Here, we use the evolutionary radiation of noctilionoid bats, the most diverse clade of mammals in terms of diet type, as a model system to show how the tooth developmental program evolved during the adaptation to new diet types. We combined morphological, developmental, cellular, and modeling approaches, to investigate the developmental differences between two tooth classes, molars and premolars and the mechanisms driving their diversification. We demonstrate that tooth classes develop through independent developmental cascades that deviate from classical models. Then we showed that the dramatic diversification of tooth number and size is driven by the modulation of the growth rate of the jaw, explaining the rapid gain/loss of teeth during the evolution of this clade. Finally, we propose a mathematical model that recapitulates the successive appearance of tooth buds and supports the hypothesis that growth acts as a key driver of the evolution of tooth number and size by tinkering with reaction/diffusion processes. Our results demonstrate developmental independence between mammalian tooth classes and provide a mechanism to explain their rapid diversification. More broadly, these results reveal how simple modifications of one developmental mechanism by another can drive the evolution of repeated structures during adaptive radiations.

## Introduction

From the conical shape of the earliest vertebrate teeth, mammals have evolved a heterodont dentition with four tooth classes (incisors, canines, premolars, and molars), each with distinct morphologies that allow for specific functions during food processing ^1^. This innovation enabled mammals’ evolution of complex teeth with an astonishing diversity of morphologies and their subsequent utilization of a broad range of dietary sources; it is therefore considered a key event in the evolutionary success of the group ^1–3^. In the last 30 years, the study of vertebrate tooth development has led to impressive new insights regarding the evolution of teeth in various clades ^2,4–6^. However, our understanding of the origin of mammalian tooth classes remains limited, in large part because most developmental studies on mammalian teeth have focused on mice. With their derived, reduced dentition containing only molars and extremely modified ever-growing incisors, mice make a less than ideal model system for studying the origins of mammalian tooth classes. The field of evo-devo therefore needs a mammalian model with a complete dentition with which to study the developmental foundation of the evolution and diversification of tooth classes.

The ideal mammalian group to fill this gap would possess a complete dentition (e.g., all four tooth classes), tremendous morphological variation in this dentition, and accessible development. Noctilionoid bats meet all of these requirements. Emerging forty-five million years ago, noctilionoid bats underwent a major evolutionary radiation such that today their more than 200 species utilize nearly all possible mammalian diets (i.e., fruit, nectar and pollen, leaves, seeds, arthropods, small vertebrates, fish, and even blood) ^7^ (Fig. 1, Extended data Fig.1 and 2). Similar to the evolution of disparate, diet-related beak shapes in Darwin’s finches ^8^, noctilionoid bats have evolved a wide diversity of skull shapes to meet their dietary needs ^7^ along with an exceptional diversity in the proportion, size, shape, and number of teeth, particularly those used for mastication (premolars and molars; Fig. 1, Extended data Fig.1 and 2) ^9–12^. In addition, tooth proportions and shape are highly variable, reflecting the evolutionary history of the clade. In association with this variation in tooth number and size, rostrum and jaw length are highly diverse, and range from extremely elongated in nectarivorous species, to highly shortened in durophagous frugivores ^13^. This last aspect of cranial morphological diversity in particular has been associated with variation in cell division rates and heterochronies during development ^11,14,15^. Together these traits make noctilionoid bats an ideal model with which to study the diversification of the patterning of mammalian tooth classes during evolutionary radiations.

**Fig. 1:**
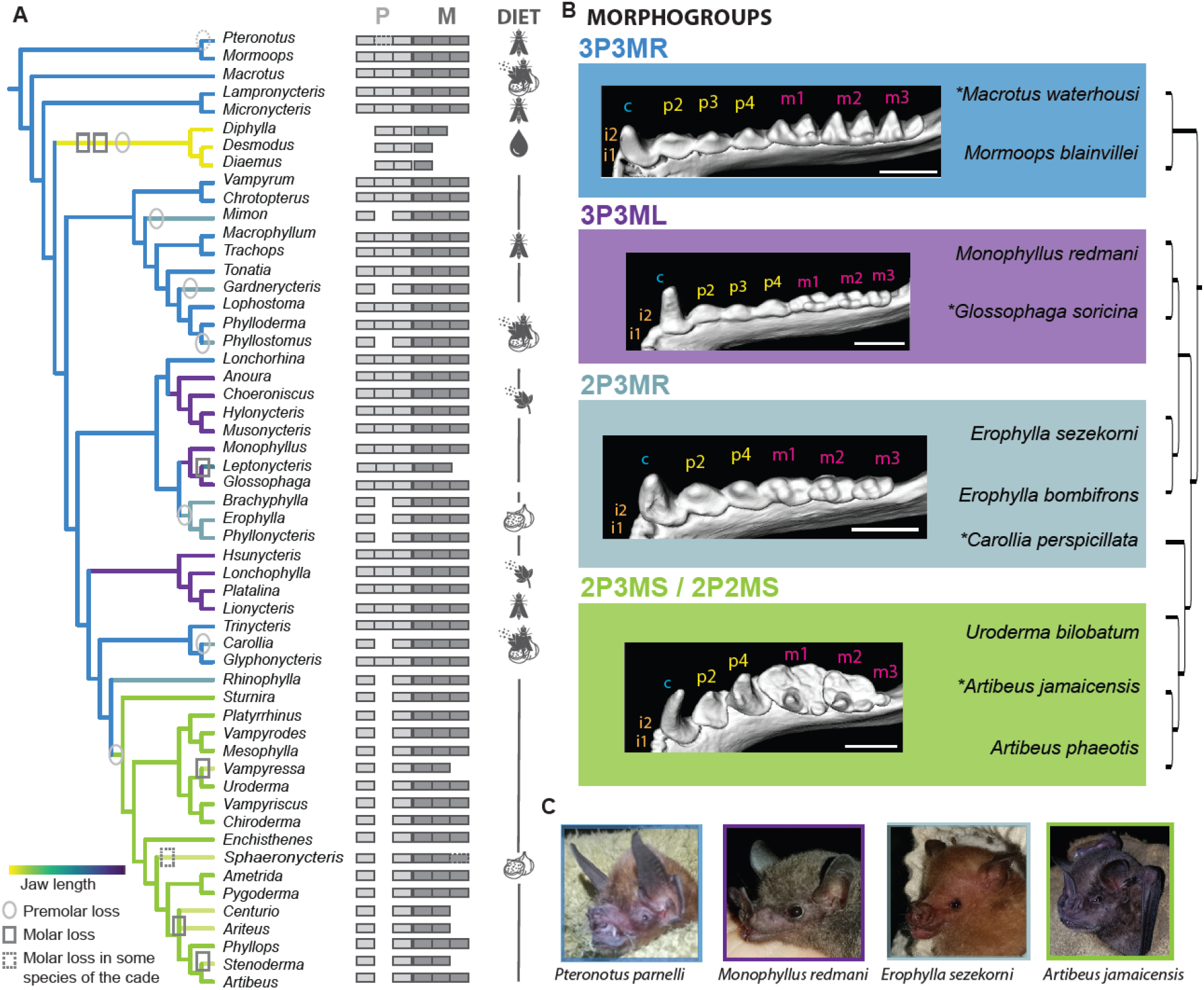
Dentition diversity in noctilionoid bats. **A**, Tooth formula and jaw length of noctilionoid bats. Jaw length is represented with a color code and premolar and molars losses events by circles or squares respectively. Some losses happened independently in different clades. Diet is indicated with icons. **B**, Morphogroups used in this study. Tooth classes are indicated by a letter: i, incisor; c, canine; p, premolar; m, molar. Representative genera and species investigated during development. **C**, Picture of four species in the four different groups representative of jaw size and tooth composition diversity.

Fifteen years ago, a groundbreaking study by Kavanagh and colleagues uncovered a developmental inhibitory cascade by which at least one class of teeth, the molars, are patterned ^16^. Through investigations of molar development in mice, this study revealed that molar development follows a simple reaction/diffusion (Turing pattern) based rule that controls the successive development of molars and their respective proportions as the dental lamina grows. However, while later research confirmed that this rule sometimes can predict the proportions of molars in other clades ^17,18^, consistent with the same developmental mechanisms being in play during their formation, nothing is known about how and if this rules also patterns the development of tooth classes beyond molars. Furthermore, studies in some mammalian clades have revealed instances in which molars explore areas of morphospace beyond those predicted by the cascade. This suggests that the cascade itself evolves and/or interacts with other developmental mechanisms to produce the tremendous variation of tooth morphologies seen in nature ^19–22^. Together, these observations highlight a need to examine the developmental cascades regulating tooth development in diverse clades to understand the developmental foundations of tooth class patterning and diversification.

In this study, we use the outstanding variation of noctilionoid teeth as a natural experiment to explore how the developmental cascades that pattern serial structures have evolved to produce tooth classes and adaptive morphological variation. We find that noctilionoid premolars and molars are formed by distinct signaling cascades and that the distribution of tooth number and size differs in the premolars and molars of extant noctilionoids. In addition, by investigating the developmental processes driving these differences, we find results consistent with the hypothesis that growth, by perturbing the underlying Turing processes, modulates the number and size of the different classes of the teeth of noctilionoid bats and possibly other ectodermal appendages in bats and other mammals.

## Results

### Noctilionoid premolars and molars deviate from expectations of the inhibitory cascade model and develop through two independent cascades

The work established by Kavanagh and colleagues has revealed that molar proportions are linked together by their developmental mechanisms^16^. To first test if two tooth classes, premolars and molars, develop through the same or distinct developmental inhibitory cascade (DIC), we calculated tooth area ratio as done in^16,23^ for successive tooth triplets in the jaw: i) the P2-P3-P4 or “premolar” triplet that (Fig. 2A), ii) the M1-M2-M3 or “molar” triplet (Fig. 2B) and iii) the P4-M1-M2 or “premolar-molar” triplet (Fig. 2C). If both tooth classes follow the same cascade, we would expect the proportions of the triplets to change in a linear manner following the relationship T(n+2)/T(n)=2(T(n+1)/T(n) -1, with T being the tooth and n being the first premolar or molar to develop ^16^ (Fig. 2D). We found that premolars and molars proportions variation are not linear and that premolars and molars proportions occupies two different morphospaces (Fig. 2A and B, Extended data Fig. 3). In addition, only 63.7% of species’ molars and 8% of species’ premolars fell within the expected IC morphospace (Fig. 2A and B). These findings suggest that the IC rule is not sufficient to predict the morphological variation observed in the premolars and molars of noctilionoid bats and that the proportions of noctilionoid premolars and molars evolve independently. To confirm these findings developmentally, we used contrast enhanced micro-computed tomography (μCT) to study tooth development in eight species of noctilionoid bats that encompass much of the diversity in tooth number and size in the clade. We found that premolars and molars form from two distinct buds (Fig. 3A and Extended data Fig. 4) that appear at the same developmental stage (Carnegie Stage 19, CS19) and seem to initiate two independent cascades (Fig. 3A) as the dental lamina grows in both directions (Fig. 3A). Together, these results suggest that the premolars and molars of noctilionoids result from two distinct developmental cascades. This finding is consistent with molar and premolar development in noctilionoids being largely independent from each other, in contrast to what has been suggested in other mammals ^23^.

**Fig. 2:**
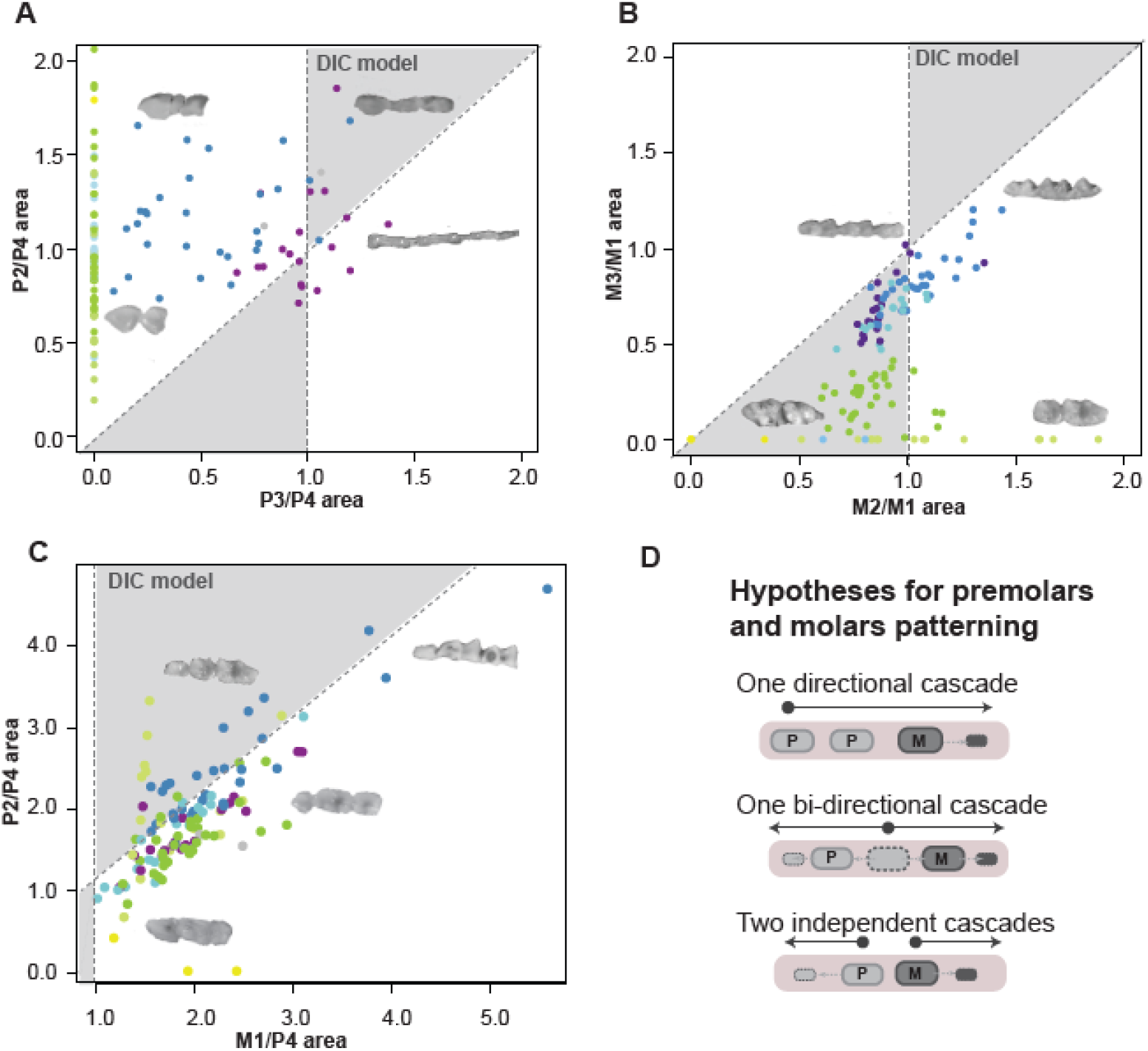
Bat premolars and molars do not follow the classical IC model. Testing the inhibitory cascade model on premolars (**A**), molars (**B**), and the P4-M1-M2 (**C**) triplet. Species that follow the IC model colonize the gray triangles. **D**, Hypotheses regarding chick-teeth development. Based on the research done in mice, premolars are molars have been hypothesized to develop through one unique inhibitory cascade that is initiated at the first premolar, here P2. Alternatively, premolars and molar could develop through one initiator tooth in both direction or through two independent cascades in opposite directions.

**Fig. 3:**
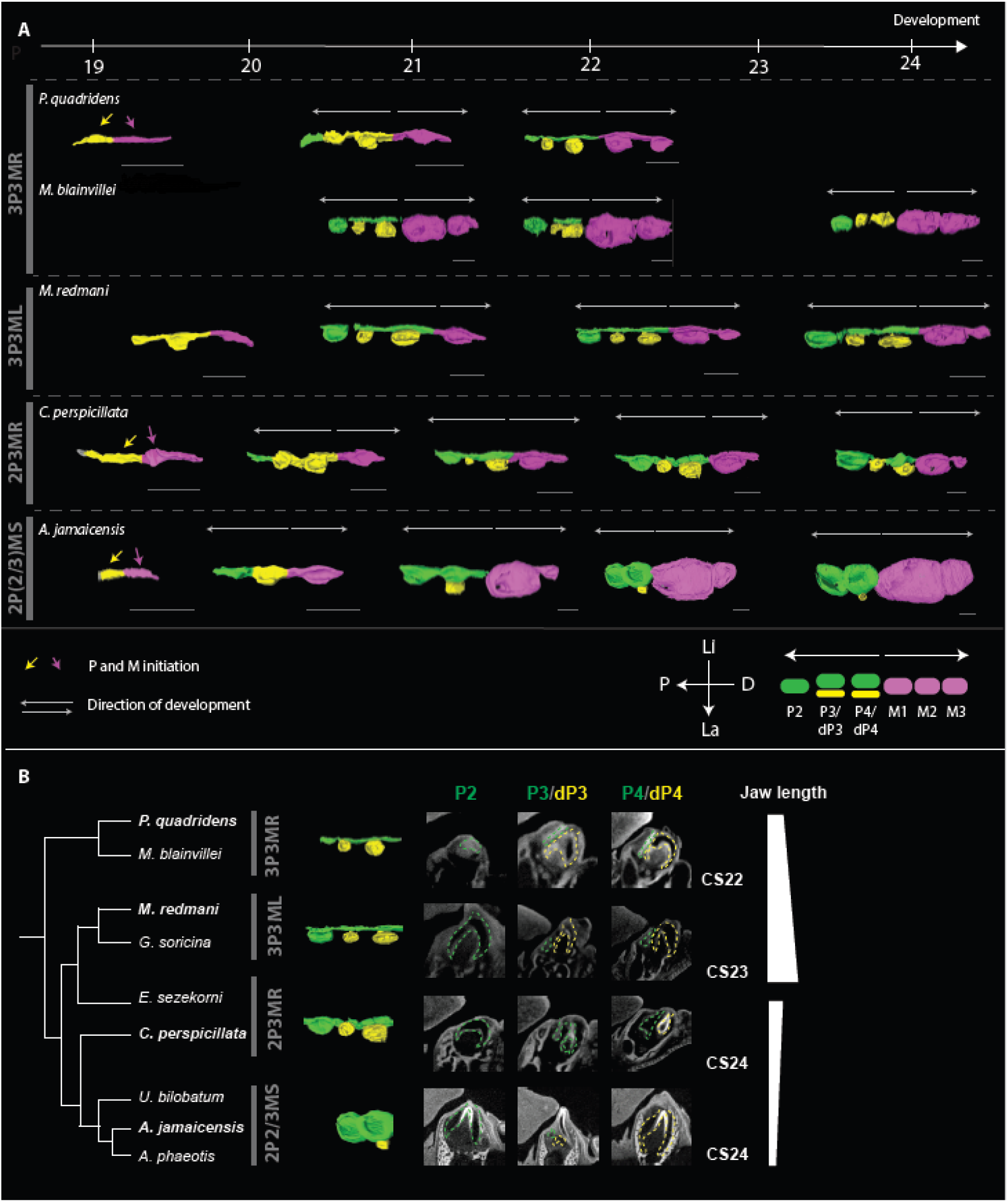
Premolars and molars develop through two different cascades and are lost differentially as jaw length decreases. **A,** Reconstruction of the developing dental lamina in our four mophogroups of bats on μCT scans. Permanent premolars are indicated in green, deciduous premolars in yellow and molars in pink. Premolars and molars develop in two different directions, as the dental lamina grows and matures. Scale bar: 200μm. **B,** Gradual loss of dP3/P3 relative to adult jaw size. μCT slides of the different premolars are represented around CS23 for the different groups. Species represented in these μCTscan are in bold, others can be found in Sup. Fig. 6. P: posterior, D: anterior, Li: Lingual, La: Labial. Scale bar: 100μm.

**Fig. 4:**
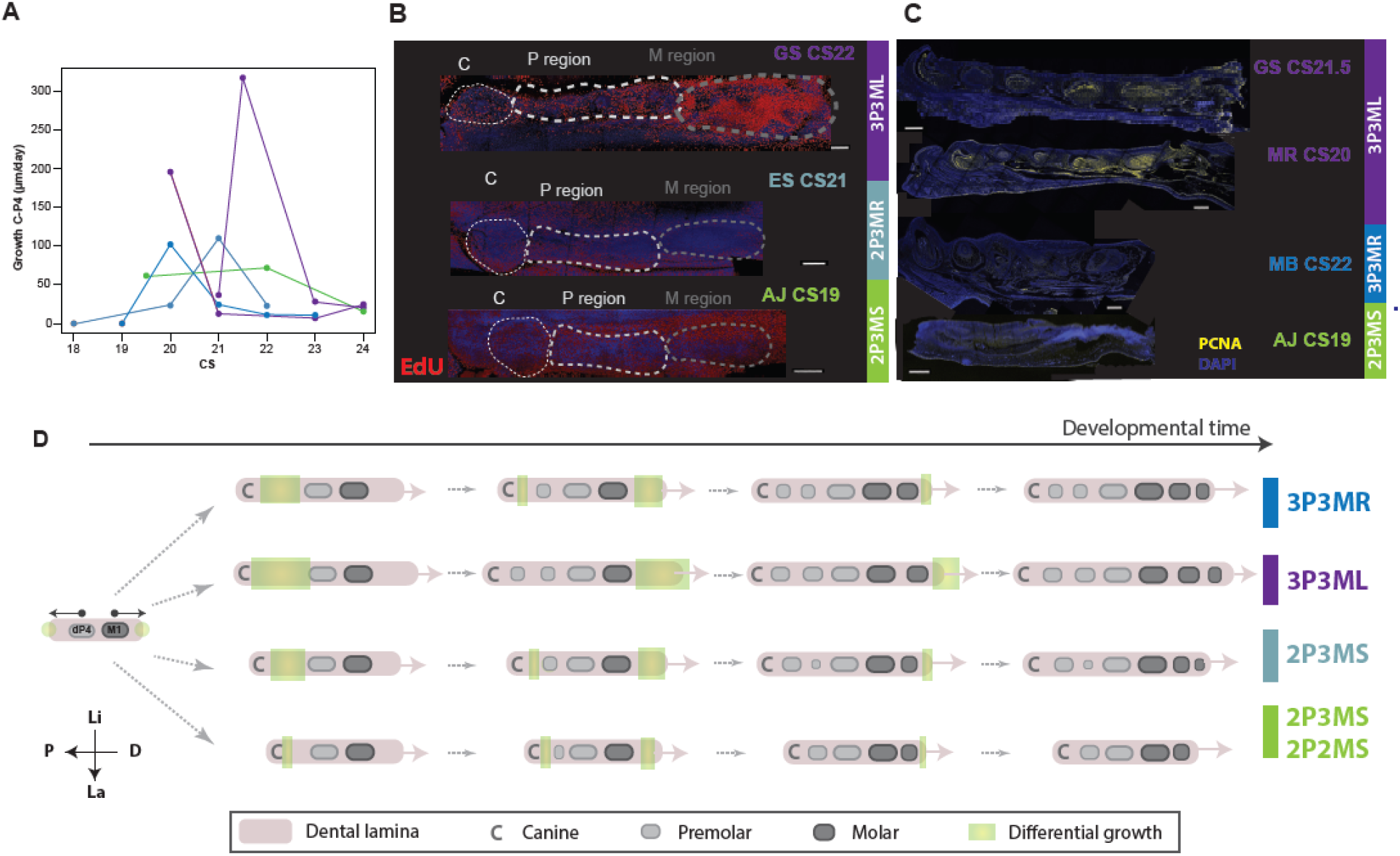
Growth variation in the premolar region. **A,** Growth rate between the Canine (C) and the P4 position during development for species in our four groups calculated from the μCT scans. **B,** Cell division in the premolar and molar regions at relevant stages for *G. soricina* (GS, 3P3ML), *M. redmani* (MR, 3P3ML), *M. blainvillei* (MB, 3P3MR), *E. sezekorni* (ES, 2P3MR) and *A. jamaicensis* (AJ, 2P3MS) labeled by EdU in whole mount dissected jaw or **C,** PCNA in sections. The position of the canine is indicated with a white dotted line. CS: Carnegie Stage. Scale bar: 200μm **D,** Dynamics of apparition of premolar and molar buds and differential growth, based on μCT scan measurements and cell division experiments. Dental lamina develops in both directions from the initial dP4 and M1 tooth buds independently for premolars and molars. Differential growth rate for the different groups is indicated in green.

**Fig. 5:**
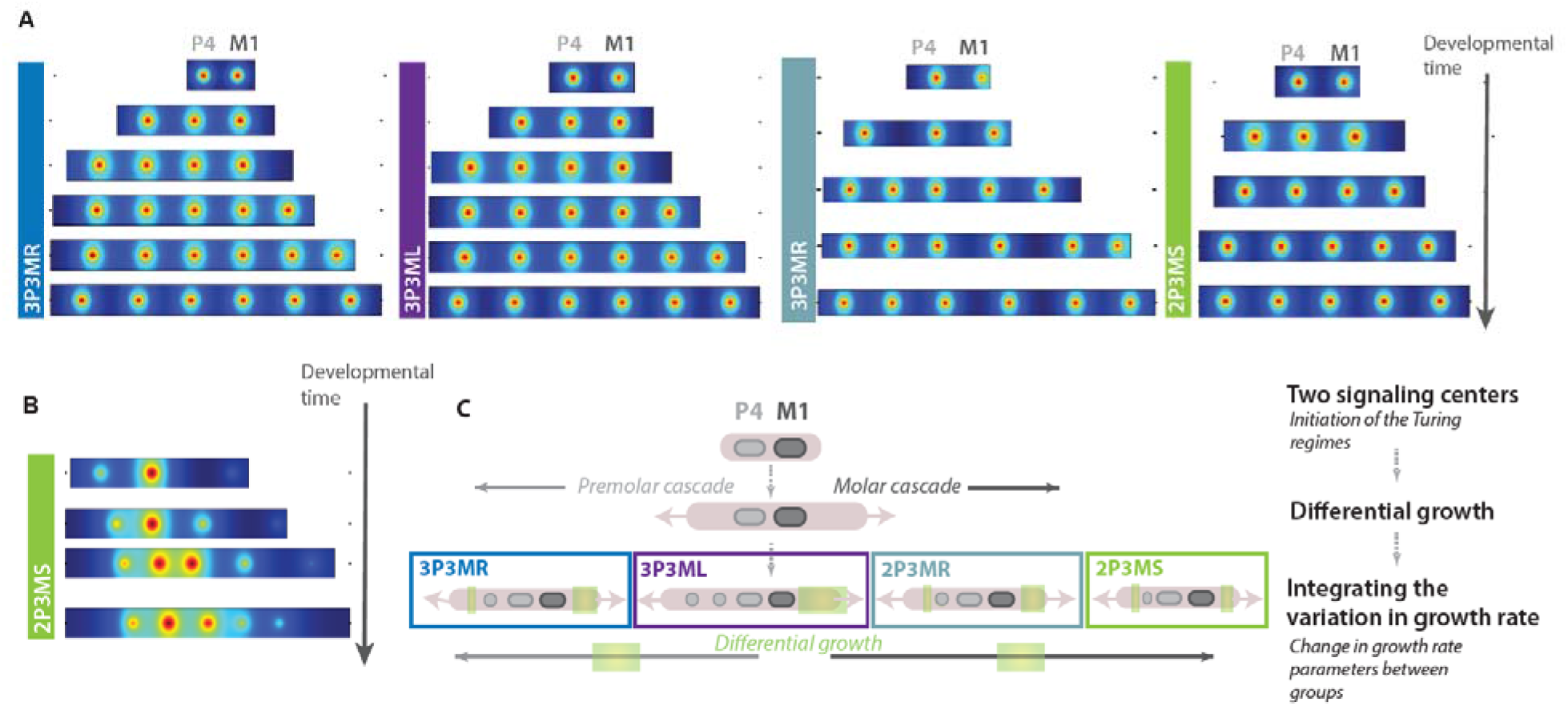
Modeling growth rate in different species is sufficient to recapitulate the variation in tooth number and size observed in Noctilionid bats. **A,** Reconstruction of the developmental sequences for 3P3MR, 3P3ML, 2P3MR and 2P3MS bats using Turing patterns incorporating differential growth. Inhibitory fields are represented by the circles. **B,** Reconstruction of the developmental sequence of *Artibeus jamaicensis* and its tooth size variation using Turing patterns with differential growth. Inhibitory fields are represented by the circles. **C,** Model: premolar and molar develop through two independent cascades, each of them having a growing domain. Variation in growth rate between the different species is integrated into the model, reproducing the diversity seen in the different morphogroups.

**Fig. 6:**
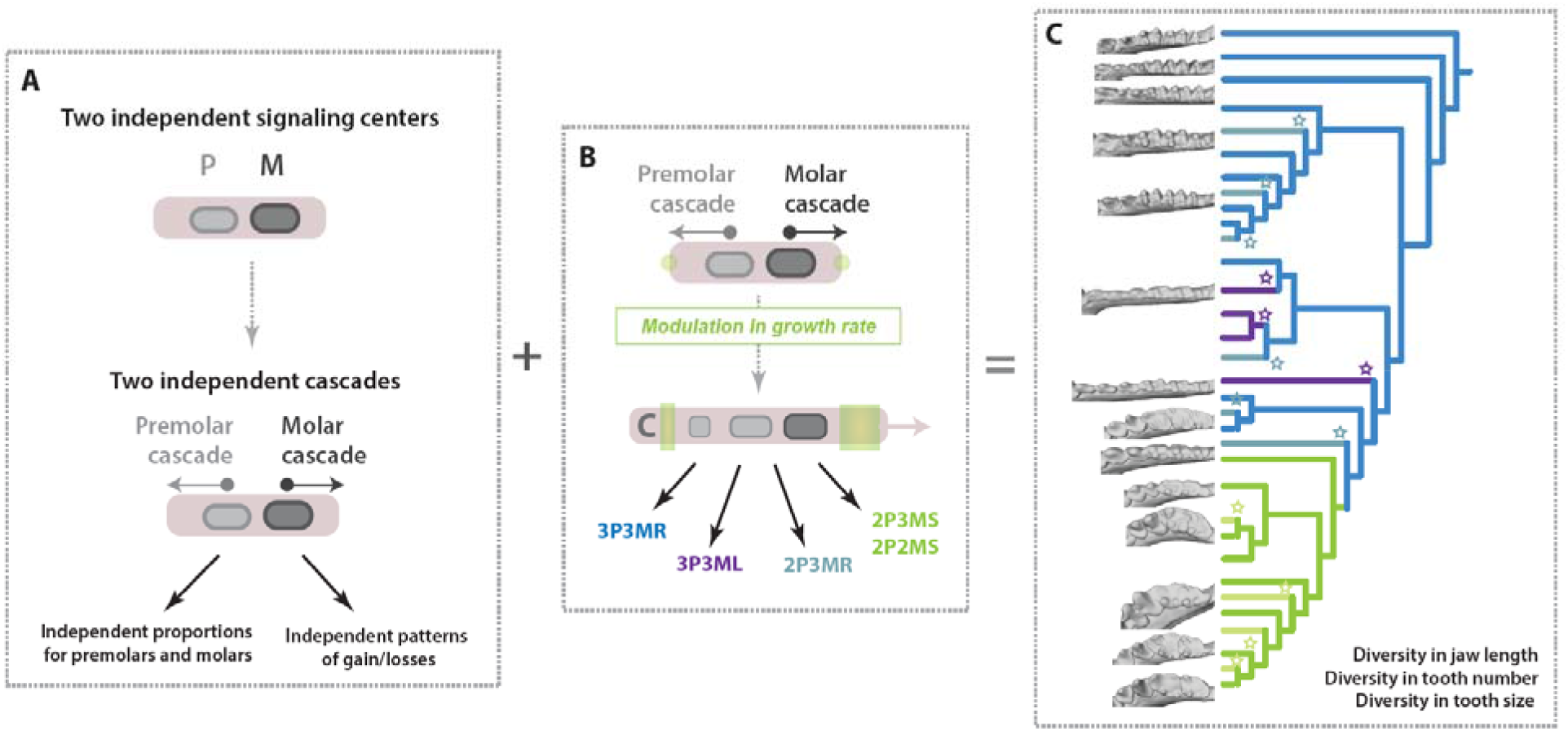
Variation of developmental cascades explain the differences between premolars and molars and their evolution. **A,** Premolar and molar develop through two independent cascades that each have their growing domain. **B,** Variation in growth rate explain the different morphogroups and losses observed in the different bats in relationship with jaw length. **C**, Variation of both the IC cascade and growth rate explain the dramatic diversity of premolar and molar number and size observed in Noctilionoid bats as well as how same morphologies have been reached convergently (stars).

### Premolar and molar numbers and proportions are tied to jaw length in noctilionoids

As tooth proportions are controlled by the underlying developmental mechanisms that regulate their formation ^16^, we quantified morphological variation in the premolars and molars of 118 species (N ≤ 3) of noctilionoid bats that span the ecological and dietary guilds found in the clade. Species were classified into four main morphogroups (see Extended data regarding other morphogroups, Extended data Fig 1, 2, 3 and 5, Extended data Table 1 Supplementary Tables 1, 2 and 3) based on relative tooth number and jaw length (Fig. 1, Extended data Fig 1): 3P3MR, the ancestral pattern, composed of insectivores and omnivores with regular jaw lengths, 3P3ML, consisting of nectarivores that have converged on an elongated jaw phenotype, 2P3MR, a convergent phenotype found in insectivores, omnivores, and frugivores with jaws of a regular length, and 2P3MS/2P2MS, containing derived, frugivorous bats with short jaws. The naming convention works as follows, a bat with 3 premolar teeth (3P), 3 molar teeth (3M), and a relatively long jaw (L) would be coded as 3P3ML, with the same teeth and a relatively short jaw (S) as 3P3MS, and with the same teeth and an average jaw length (R) as 3P3MR. We found the number of teeth to generally be associated with the size of the jaw among species (Extended data Fig. 5); the jaws of 6-toothed 3P3ML and 3P3MR bats are significantly longer than those of 4- of 5- toothed 2P3MR, 2P3MS and 2P2MS bats. Interestingly, some of these morphogroups, such as 2P3MR or 2P2MS, are polyphyletic and contain independent events of tooth loss, consistent with jaw length reduction having been being repeatedly associated with tooth loss in noctilionoids. In addition, we found that the proportions of individual premolars and molars are more disparate in species with shorter jaws (typically frugivores; Extended data Fig. 3); bats with elongated jaws routinely have thinner and longer teeth with more similar proportions (3P3ML, Extended data 2 and 3, Fig. 1). These findings are consistent with an association between the processes of jaw elongation and the generation of tooth proportion and number.

### Premolar and molar patterning and gain/loss is associated with dental lamina length

To explore the developmental mechanisms linking tooth number, tooth size, and jaw length, we investigated the possibility that the sequence of premolar and molar appearance during development depends on the available dental lamina space. As the dental lamina develops, the teeth appear successively so the space available at a given developmental time for the teeth depends on the growth rate of the dental lamina. In 3P3MR bats, we observe that development of dP4 is followed by the sequential development of dP3 and P2 (Fig. 3A and B) which is consistent with the predictions of the classic IC model and what is seen in mice molars, albeit in the reverse direction (i.e., from the back to the front of the jaw) (Fig. 2C). In 2P3MR bats, which have slightly shorter jaws, further examination of the developing dental lamina reveals that, while the dP3 forms and mineralizes, its replacement adult P3 is initiated but does not develop further (Fig. 3B). At the extreme, in 3P3ML bats that exhibit an elongated jaw, dP3 and P2 appear almost simultaneously as the jaw develops rapidly (Fig. 3A and B). Together, these results suggest that the development and timing of formation of different premolar buds is influenced by the space available in the jaw, likely because new teeth are able to form only at a certain distance from each other as predicted by the IC ^16^. This finding is supported by the pattern of tooth formation in shorter-jawed 2P2MS and 2P3MS bats; the development of the dP3 is initiated but the incipient tooth fails to grow and/or mineralize resulting in the loss of both dP3 and P3 (Fig. 3B and Extended data Fig. 5). In 2P2MS bats, such as *A. phaeotis*, the M3 is lost with no evidence that it ever started to develop (Extended data Fig. 5). These patterns of loss are consistent with the variation observed in adult bats: the smallest teeth, specifically the middle premolar (P3) and the last molar (M3), exhibit the most variation in size among species with short jaws (Extended data Fig. 3); P2 and P4 and M1 and M2, respectively, exhibit similar size variability among species with short and elongated jaws. Of note, the loss of the M3 is polymorphic in some species of short-faced bats, and this within-species variation has been linked to subtle variation in jaw size among individual bats ^24,25^. These observations suggest that the presence/absence of M3 is dependent on the available space in the developing jaw (e.g., *A. watsoni* appears to be just at the limit condition for which the M3 does or does not develop further). To sum up, premolars and molars exhibit divergent patterns of loss with decreasing jaw lengths, with premolars losing the middle tooth gradually (dP3 and then P3) and molars the last tooth of the row, the M3. These losses appear to have happened convergently in bats with similar jaw lengths.

### The number and proportion of premolars and molars is associated with variation in growth rate

The results presented above support the hypothesis that the numbers and proportions of premolars and molars are linked to the length of the jaw and, in particular, how fast the jaw is growing and where that growth is located along the jaw at the time of tooth bud formation. To further explore this idea, we measured the growth rate of the premolar area during development in 3D models reconstructed from phosophotungstic acid-contrasted μCT scans in eight focal species representatives of our four morphogroups (see methods). These measures reveal that species with jaws of average length (3P3MR and 2P3MR) exhibit a moderate peak of jaw growth around stage 20 as dP3 and P2 develop (Fig. 4A). In 3P3ML bats with elongated jaws, this peak is 3 times faster than that of average jaws and corresponds to the almost simultaneous formation of dP3 and P2. In contrast, no growth peak was observed in short-faced bats (2P3MS and 2P2MS); growth rate was lower in short-faced than other bats in this region. To examine these patterns in more detail, we used EdU and PCNA labeling to trace cell proliferation during jaw development (Fig. 4B and C, Extended data Fig. 6). We found that 3P3ML bats seem to exhibit higher rates of cell division in the premolar area compared to species with other morphotypes, while 2P3MS and 2P2MS bats tend to show the lowest rates of cell division. These results echo previous results in long-jawed nectarivorous noctilionoids ^14^ and support the idea that differences in cell proliferation or growth rate contribute to and possibly drive craniofacial ^15^ and tooth size ^18^ differences between species (Fig. 4D).

### Growth rate likely tinkers with Turing patterns and thereby modulates the appearance order and size of teeth

Our findings suggest that the growth rate of the jaw might perturb both the sequence of tooth appearance and the reaction/diffusion mechanisms that shape tooth proportions. As these two parameters depend on the activation/inhibition processes that pattern repeated structures, we postulate that the variation in tooth number and size observed in noctilionoid bats could possibly be simply explained by growth rate-induced perturbations on the Turing mechanisms behind the ICs. To begin to test this, we computationally examined if we could reproduce the various phenotypes observed during development by modeling variation in jaw size and growth. We implemented a simple model of a Turing-type reaction-diffusion system (Supplementary Methods 1) to recapitulate the different sequences of insertions observed for the two independent cascades for premolars and molars (Fig. 5 A and B). Using only apical growth, we successfully recapitulated two of the four morphogroups (corresponding to 3P3MR *P. quadridens* and 3P3ML *M. redmani*) (Fig. 5A). Presupposing exogenous spatial gradients (plausibly corresponding to a pre-pattern or additional spatial modulation), we also captured the insertion sequence and –qualitatively– the size variations observed in 2P3MS *A. jamaicensis* (Fig. 5B). These results demonstrate that, while there are undoubtedly other factors influencing the insertion and modulation of tooth signaling centers (which could explain the 3P2M group that loses a molar while having a long jaw with diastemas; Extended data Fig. 2, 3 and 5), simple models combining reaction-diffusion processes and differential growth are sufficient to explain much of the observed variation in noctilionoid teeth. This supports the importance of growth rate and jaw length variation in modulating the number and size of both premolars and molars in two distinct ways (Fig. 6).

## Discussion

Until now, the developmental processes driving the differences among mammalian tooth classes have remained relatively obscure ^5^. From a morphological point of view, the differences between tooth classes have often been assessed through morphological modules ^26–28^ or through the inhibitory cascade model ^23,26^. These methods commonly identify modules specific to the incisors and canines but fail to identify distinct premolar and molar modules, possibly because they lack the precision to distinguish these classes and/or the morphological and perhaps genetic differences between these latter tooth types are less pronounced than for other classes. On the other hand, developmental studies, largely based on morphological observations of the dental lamina of species ^29^ including the ferret ^30^, shrew ^31,32^, straw-colored fruit bat ^33^. opossum (^34^ and personal observations), and this study reveal that lower premolars and molars tend to develop in opposite directions, and thereby support the hypothesis that premolars and molars are patterned through independent mechanisms. From a molecular perspective, some evidence suggests that the early patterning of the jaw is driven by a “homeobox-code” that divides the jaw into territories that determine tooth classes prior to bud development (reviewed in ^35–37^). While these results suggest that tooth class specification and/or determination might occur prior to placode induction, results of other studies suggest that this early code is not sufficient to establish tooth class identity. In line with this, recent studies in the lizard genus *Pogona* have demonstrated that a heterodont dentition can be achieved through a simple modulation of Eda signaling during later tooth development^38^. In opossum, shifts in gene expression patterns have been observed between tooth classes, suggesting that different core developmental programs control their formation^34^. Together, these results suggest that events that occur during later tooth formation and morphogenesis (e.g., bud, cap and bell stages) likely play a major role in the establishment of tooth identity. The work we present here, which suggests that premolars and molars develop independently by two different cascades, supports the hypothesis that the development of at least the premolar and molar tooth classes are largely independent and uncovers a mechanism that could act in tandem with the homeobox code and other processes to modulate that development.

Rapid diversification in the number and size of teeth is common during the colonization of different ecological niches and the associated incorporation of new food sources into the diet ^1,39–41^. However, little is known about how this diversification could be facilitated by the developmental mechanisms controlling tooth development. In particular, molar development and proportions have been shown to be constrained in mammals through the IC ^16–18,42^, making it traditionally difficult to explain the colonization of new morphospaces. While the expansion of studies to include more clades has revealed that some seemingly do not follow the expectations of the IC model ^19,21,22,42^, the mechanisms by which these species escape this developmental bias remain unclear. Here, we provide some possible resolution to this conundrum by showing that growth, by perturbing the IC during tooth patterning, could act as a simple modulator of tooth number and size and push tooth proportions into new areas of morphospace that are not predicted by the IC itself. In addition, we show that premolars and molars likely develop through two different cascades. This finding helps explain observed differences between premolars and molars in basic morphology and evolutionary changes in that morphology (e.g., proportion, loss) coincident with the adoption of new food sources.

In our study, we showed results consistent with the hypothesis that growth rate and the resultant jaw space, by modulating Turing patterns, rapidly modulated the number and size of the teeth of noctilionoid bats during their evolutionary radiation (Fig. 6). This finding is in accordance with the patterns of tooth appearance in the upper jaw of shrews ^31,32^. In shrews, premolars appear antero-posteriorly (while premolars develop postero-anteriorly in the lower jaw) with P3 appearing before P4, suggesting that the first tooth from each class to appear could initiate its own cascade whose direction will depend on the space available in the jaw ^5,32^. This could also explain how the direction of the addition of new teeth could change between species. Beyond teeth, this finding is consistent with what has been suggested for other ectodermal appendages (e.g., palatal rugae in mice and hamsters), in which a growth burst drives the appearance of new segments in the resulting available space ^43,44^, and in other ectodermal organs ^45,46^ that follow Turing pattern formation but without directional growth. As both the IC cascade and growth are implicated in the development of ectodermal appendages in general ^16,17,47,48^, our conclusions are potentially applicable to other ectodermal organs and propose a testable model to explain how their number and proportions can rapidly evolve, simply by modulating growth rate.

## Conclusion

Our work demonstrates how new morphologies can potentially be rapidly achieved through subtle changes in the interaction between multiple developmental biases during the bursts of diversification that often accompany evolutionary radiations (Fig. 6). While studies of developmental rules often solely focus on the evolution of one developmental process to explain the evolution of characters, our work reveals the importance of studying the complex interaction of different developmental processes to fully understand the evolution of new morphologies during the colonization of new ecological niches, and identify bat teeth as a new model system to study these questions. Further work should focus on the identification of the molecular basis of these processes and how they interact with other mechanisms.

## Online methods

### Bat groups

We grouped species in 6 morphogroups representative of their dental diversity (Extended data Fig 2, Supplementary Table 2) relative to tooth number and jaw size. 3P3MR represents the ancestral pattern of 3 premolars and 3 molars and a regular jaw length. 3P3ML represents bats with 3 premolars and 3 molars with elongated jaws as seen mostly in nectar feeders. 2P3MR represents bats with 2 premolars and 3 molars with a shorter jaw length and 2P3MS and 2P2MS represents bats with 2 premolars and 2 or 3 molars, with a short face and jaw. The M3, present in 2P3MS is extremely reduced. In our analyses, because the size of the jaw is an important feature of short-faced stenodermatinea bats, we grouped 2P2MS and 2P3MS bats. In our data set, we also found two species with three premolars and two molars, 3P2M. In these species, the P3 is extremely reduced and their jaw size is closer to the second group. Because this morphology is marginal, it has not been used for the following analyses but was kept for the modeling. Finally, we excluded the vampire bats from our analysis given the lack of embryos and their extremely reduced and derived dentition (2P1M or 2P2M) that limit our developmental investigations. Diet groups were assessed based on ^13^.

### Museum and Field specimens

Museum specimen pictures have been: 1) taken from the FMNH in Chicago with a Nikon camera, 2) downloaded from Animal Diversity Web https://animaldiversity.org/ (see Extended Data Fig. 2). Field specimens were collected in the field (see Supplementary Table 1) using mist nets, harp traps or butterfly nets and euthanized humanely with isoflurane according to approved institutional animal care and use committee (IACUC) protocols 14199 at UIUC, 2017-093 at UCLA and the following permits Dominican Republic: VAPB-01436; Puerto Rico: 2015-EPE-028; Trinidad: 000619 and 000620 Apr 18th 2018. Specimens were then fixed ON at 4C with PFA and dehydrated the next day in 100% methanol and stored at -20C until used.

### Body mass

Body mass (Supplementary Table 2) was used to normalize the data in our analysis. Body mass data has been collected from Davalos and Melo, Chapter 8 ^7^, which gathered an impressive dataset on the body size of Noctilionoidea with the exception of: *Chilonycteris macleayii=Ptenoronus macleayii* ^49^; *Leptonycteris nivalis* ^50^; *Lonchophylla thomasii* (Emmons, 1990); *Platyrrhinus fusciventris* ^51^.

### Statistical analysis

Differences between the tooth areas of the different groups and teeth were compared by ANOVA and Tukey multiple comparisons of means (Extended Data Table 1 and Supplementary Table 3) in R.

### Developmental stages

Developmental stages for the different species of bats have been based on the development of *Carollia perspicillata* ^52^.

### μCT scanning and dental lamina segmentation

Bat embryo jaws were dissected and stained in 0.3% (Phosphotungstenic acid, Sigma) PTA in 70% ethanol (museum specimens) or 100% methanol (field samples) for 24 to 36h on a rocker at room temperature. Stained specimens were mounted in a 1.5 or 2 mL eppendorf tube between two pieces of foam and μCT scanned in a Skyscan 1172, Scanco uct50 and a Xradia BioMicroCT. Scan parameters were adjusted depending on specimen size and morphology, and voxel size ranged from 1 to 5 μm per scan. Raw μCT-scan shadow images were reconstructed to slices in NRecon, then imported into Mimics, where the dental lamina was segmented using the lasso tool every 4 to 5 slides before using the interpolation tool. Surface (stl) files were exported and used for visualization and morphological comparisons.

### Dental measurements

Adult dental measurements were taken from museum specimen photos (see Supplementary Table 1 for the specimens list) or Animal Diversity Web (animaldiversity.org) photo using ImageJ. Scale was set using the scale bar on the pictures. Crown width and length were measured three times for each tooth to ensure reproducibility. For analysis, individuals from the same species and locality were aggregated. Data available upon request. Jaw length was measured from the tip of the jaw to the middle point between the left and right angle of the jaw.

Embryo dental measurements were taken from reconstructed .stl files in mimics using the measurement tool. Each distance was measured three times, to ensure reproducibility, between the primary enamel knots or its resulting cusp to measure the distance between teeth, or the tooth primary enamel knot and the end of the dental lamina to measure tooth sizes Dental measurements are available in Supplementary Table 4.

### HCR-IHC imaging

Field sampled embryos were embedded in OCT and sectioned using a cryostat CM1520. Proliferated cells were detected using a PCNA antibody (Rabbit mAB #13110, Cell Signaling technology) at 1:300 and the signal amplified using the HCR system from Molecular Instrument according to the manufacturer protocol ^53^. Sections were imaged using a Leica SP8 confocal.

### EdU staining

Pregnant females were injected in the field with 20mg/kg of EdU reconstituted in DMSO and PBS in an intraperitoneal injection according to IUIAC procedures. 45 minutes after the injection, bats were euthanized with isoflurane. Embryos were dissected out of the female and the jaws were carefully dissected and fixed overnight in 4% PFA at 4C before being dehydrated the next day in 100% methanol and stored at -20C until imaging. In the lab, half dissected jaws were rehydrated and clarified using Scale S ^54^. After the incubation in the S3 reagent, jaws were re-hydrated in PBS before labelling. The next day was stained with a Click-iT EdU Alexa fluor 647 labelling kit according to the manufacturer protocol except for the incubation time, adjusted to 3h, at RT. After the reaction, jaws were put in Scale S4 for the final clarification and imaging at the Leica SP8 DIVE two-photon microscope at the CNSI facility at UCLA using 760nm and 1240nm wavelength for Hoescht and Alexa Fluor 647 respectively. Resulting photos were processed using LASX and ImageJ.

### Modeling

See Supplementary Methods 1.

## Data availability

The μCT scans datasets generated during and/or analyzed during the current study are available from the corresponding author on reasonable request. All other measurement datasets are included in this published article (and its supplementary information files).

## Acknowledgments

We thank A. Couzens, A. Tucker, Y. Gibert and V. Laudet for their critical reading of the manuscript. We thank L. Yin from X-ray imaging facility at UIUC, Beckman Institute for his assistance regarding CT-scanning. Two-photon excitation laser scanning microscopy was performed at the Advanced Light Microscopy/Spectroscopy Laboratory and the Leica Microsystems Center of Excellence at the California NanoSystems Institute at UCLA with funding support from NIH Shared Instrumentation Grant S10OD025017. We thank N. Simmons for museum specimen access and for organizing the field trip in Belize with B. Felton. In addition, we thank L. Rostant in Trinidad, M. Santiago and J. Almonthe in Dominican Republic and A. Rodrigues in Puerto Rico for their support in the acquisition of permits for sampling and sample exportation. We thank N. Rochette for his help with statistical analysis and M. Oliva for her comments on the results. We thank the National Science Foundation (NSF) for supporting A. Sadier and K. Sears with grant award 201780 and S. Santana with grant award 2017738.

## Author contributions

A.S. and K.S. conceived, directed the study and supervised the study. A.S., N.N., S.S., and K.S. collected the data. A.S. performed the investigations, designed the experiments and the analyses. A.K. and R.D. performed the modeling experiments. A.S, M.L. (core facility) and L.B. (core facility) performed the imaging. A.S. and R.H. processed the images. A.S. wrote the original draft and A.S, K.S, S.S, N.A. reviewed and edited the manuscript. A.S. and K.S. administrated the project and acquired funding.

## Additional information

### Supplementary Information

Supplementary information is available for this paper.

## Extended data figure/table legends

**Extended data Fig. 1:**
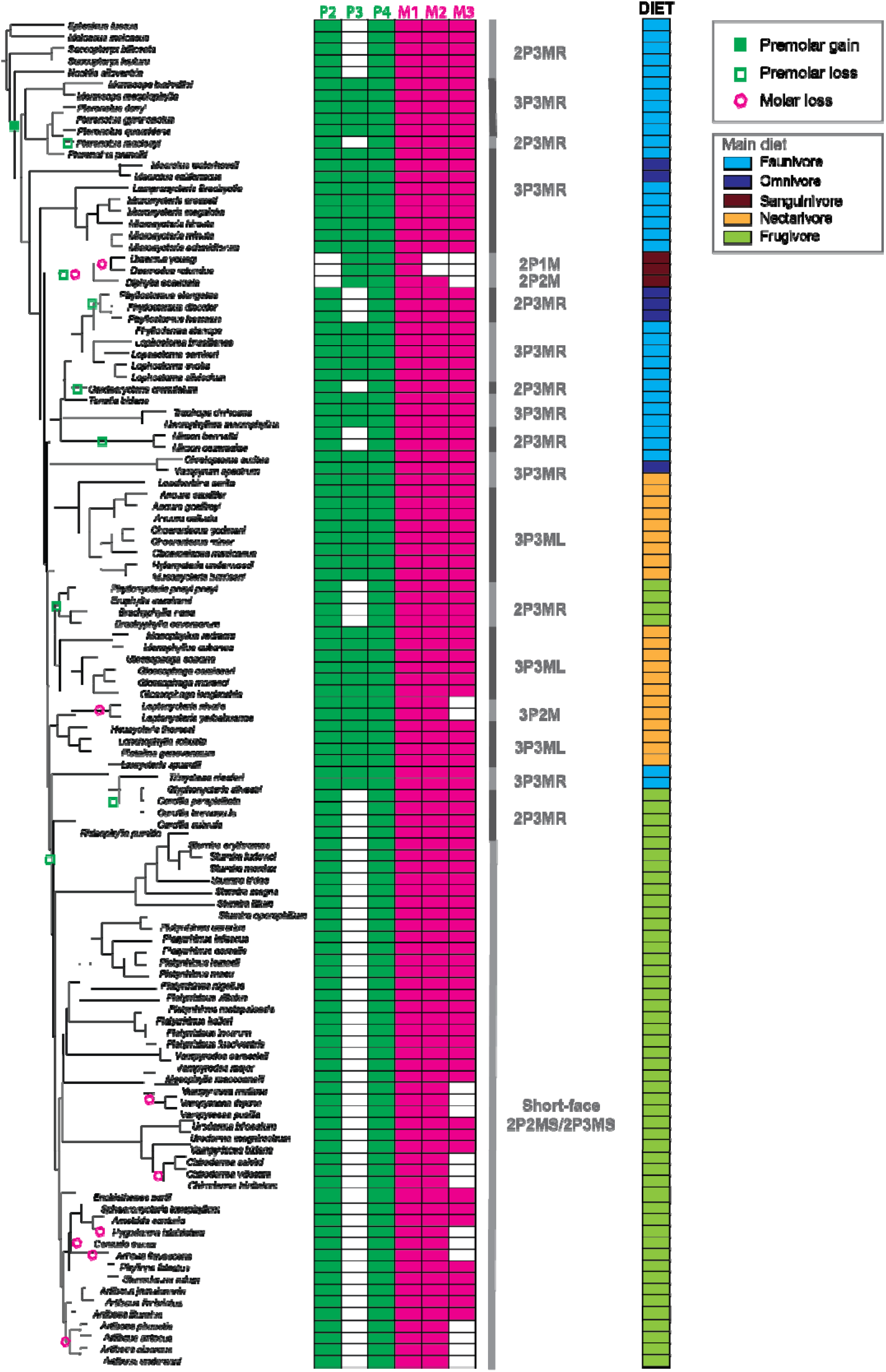
Tooth number, jaw length and diet of the different bats. Phylogeny of all the species used in our dataset, their dental formula and the seven initial groups based on jaw length and tooth number (see Extended Data Tables 1 and 3). For each species, the main diet is indicated. The events of premolar and molar gains and losses are also indicated on the tree.

**Extended data Fig. 2:**
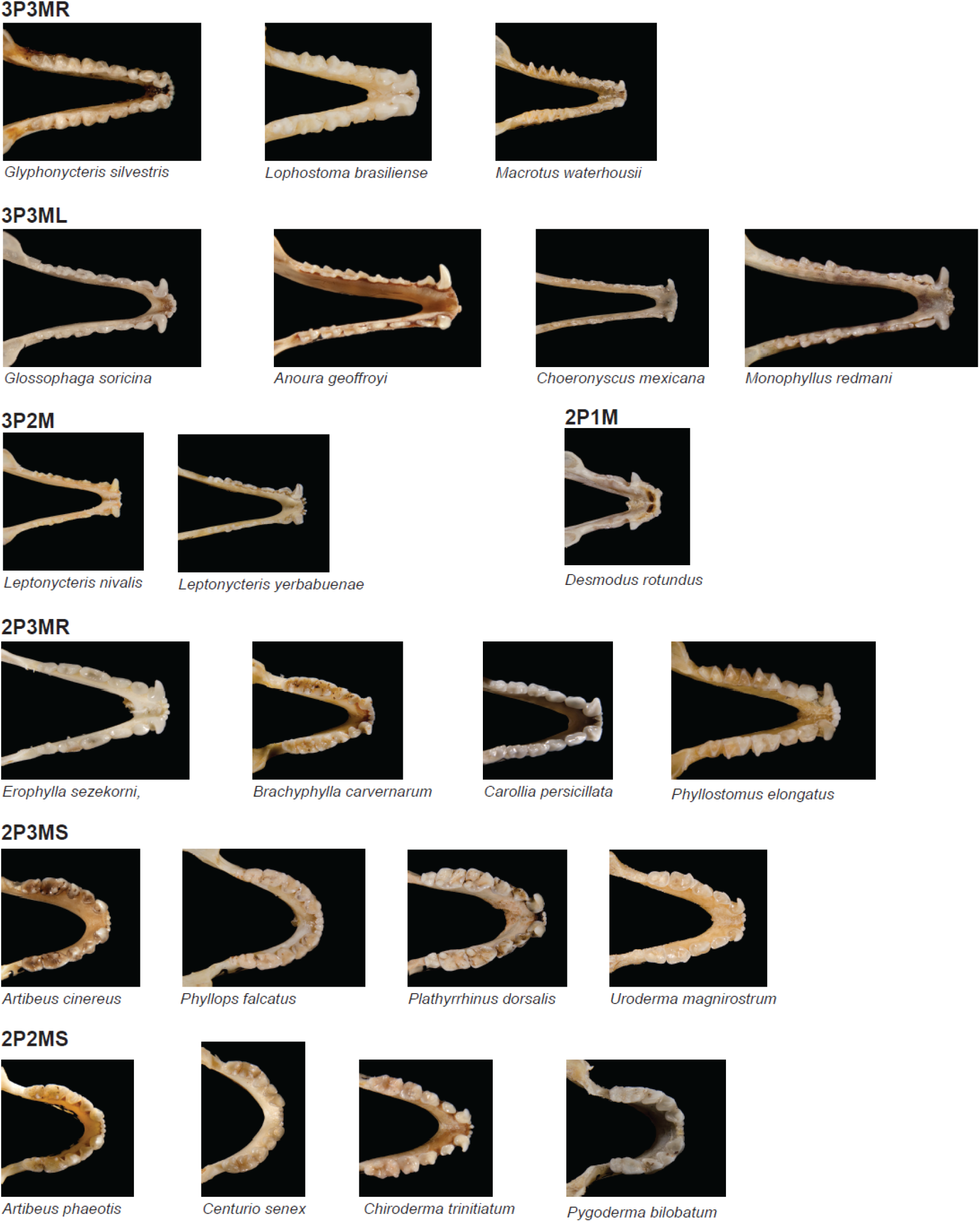
Representative bats from the different groups based of tooth number and jaw length. Pictures from ADW (animaldiversity.org, Phil Myer).

**Extended data Fig. 3:**
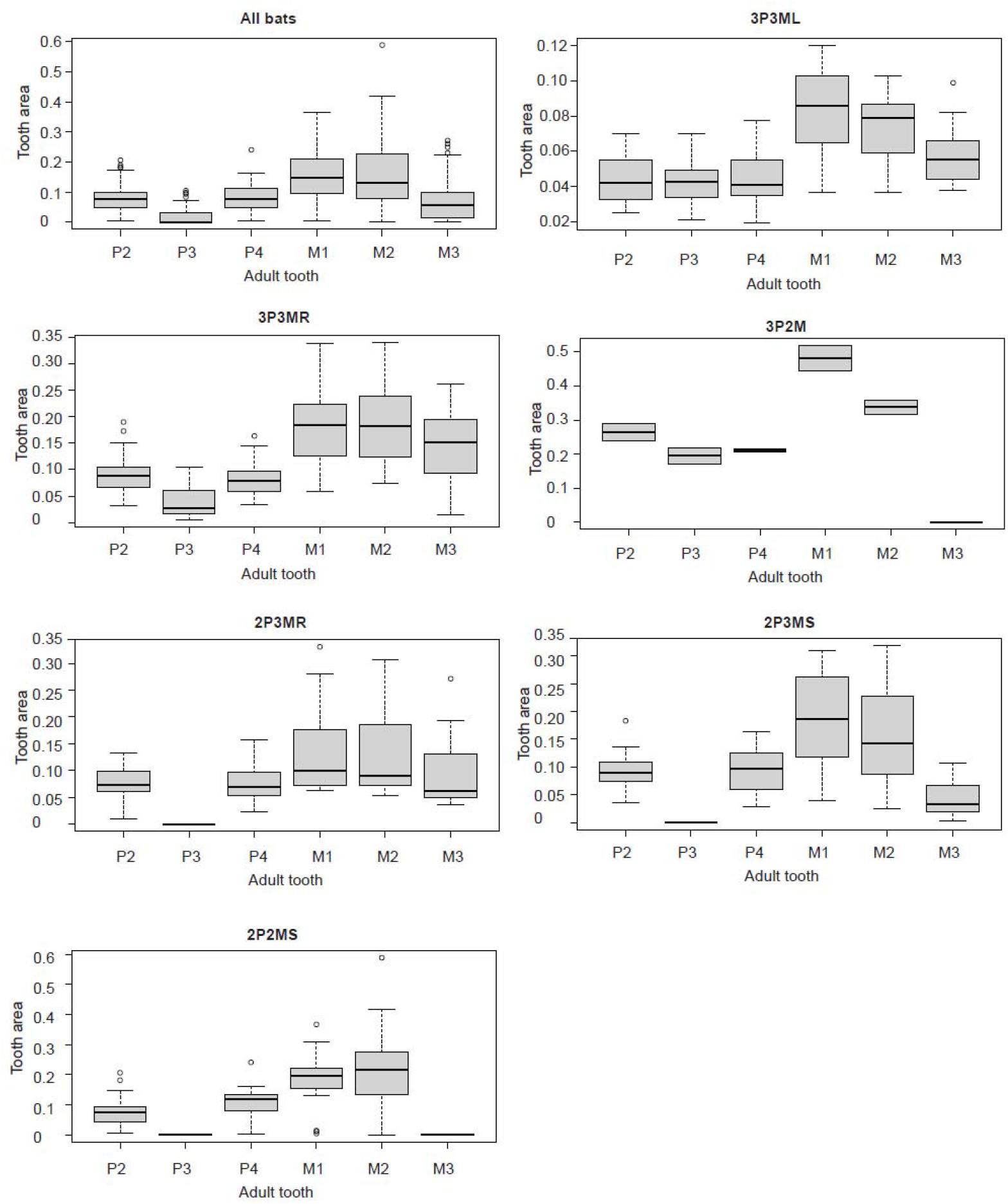
Tooth proportion variation in the different groups of bats. Tooth proportions variation in the different groups of bats (see Methods). As jaw length diminishes, molar proportions become less equal, in particular for 2P3MS. Most of the variation in terms of M1 and M2 proportions in all bats is explained by the 2P2MS group. Tooth sizes are normalized by adult body mass.

**Extended data Fig. 4:**
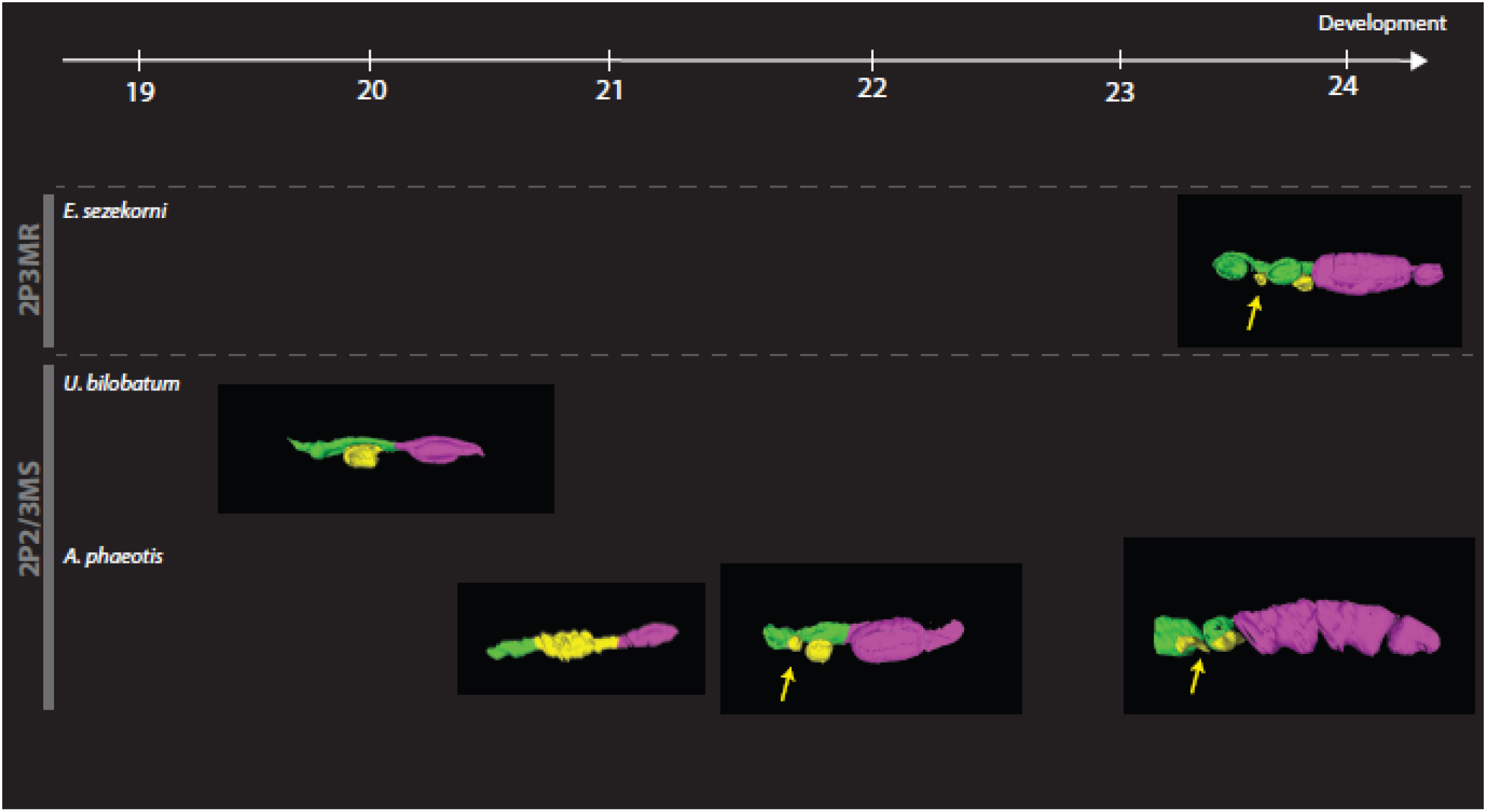
μCT scan on isolated stages for more species. Reconstruction of the developing dental lamina in three other species of bats for which the developmental series is incomplete. Permanent premolars are indicated in green, deciduous premolars in yellow and molars in pink. For *E. sezekorni*, the dP3 is indicated with a yellow arrow and will give rise to a functional dP3. For *U. bilobatum* and *A. phaeotis*, the developing dP3 is indicated with a yellow arrow at its earlier stages but then degenerates before its eruption in the juveniles.

**Extended data Fig. 5:**
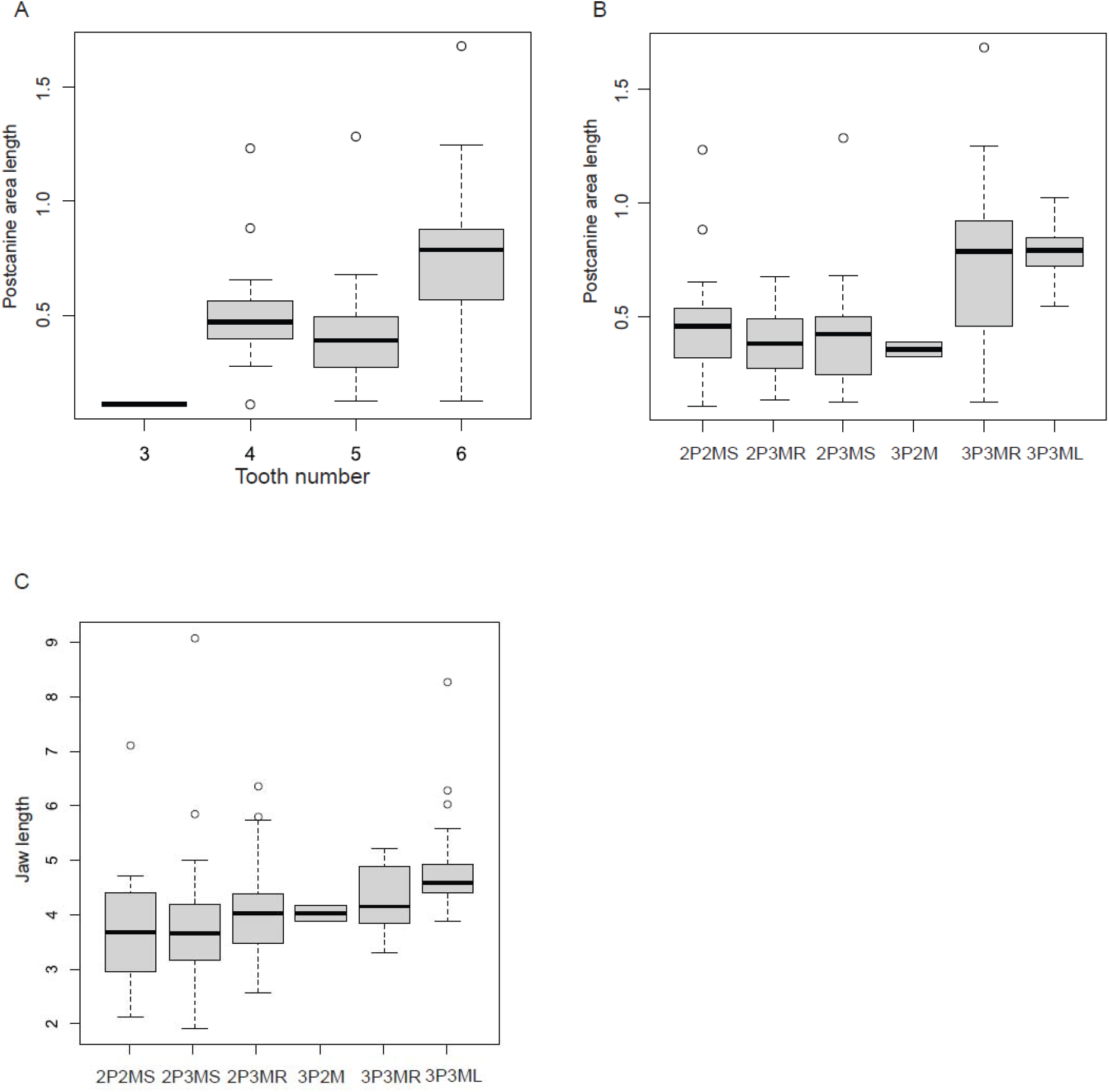
Relation between tooth number and jaw length. A: Tooth number is correlated with postcanine dental field length. B: Dental field length variation in the different groups. C: Jaw length variation in the different groups.

**Extended data Fig. 6:**
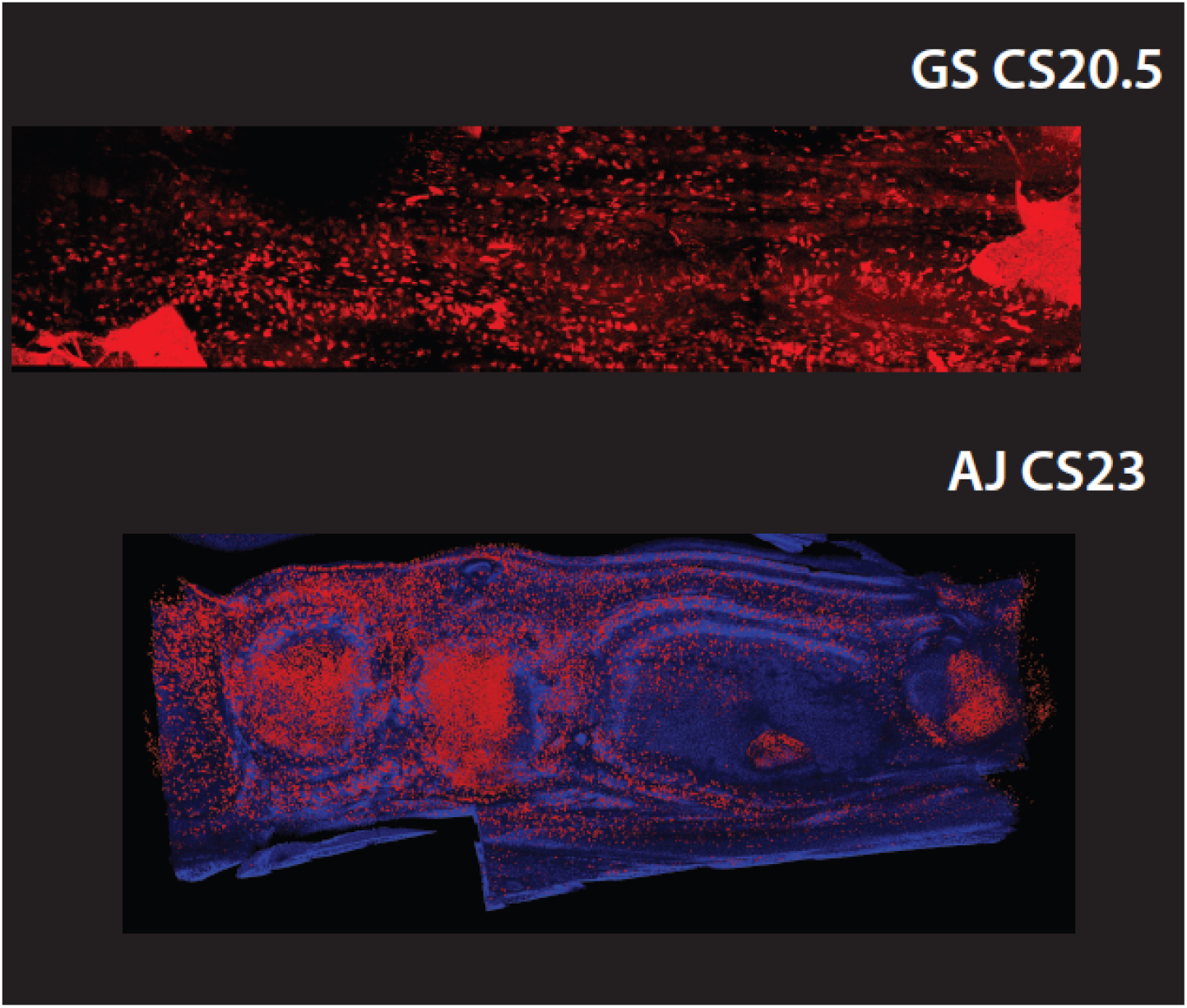
EdU from two-photon imaging. Additional species and stages (GS CS20.5: *G. soricina*, carnegie stage 20.5; AJ CS23: *A. jamaicensis*, carnegie stage 23).

**Extended data Table 1:** Size of the dental field and dental formula. A, B: Tukey multiple comparisons of means of morphogroups dental field lengths normalized by body mass (A) or P4 area (B). 2P2MS morphogroup significantly different from 3P3ML and 3P3MR. 2P3MR morphogroup is significantly different from 3P3MR and 3P3ML. 2P3MS morphogroup is significantly different from 3P3ML and 3P3MR. C, D: Tukey multiple comparisons of means of tooth number dental field lengths normalized by body mass (C) or P4 area (D). Dental field length of jaws with 4 or 5 postcanine teeth are significantly smaller than jaws with 6 teeth.

|  |  |  |  |  |
| --- | --- | --- | --- | --- |
| **A: Morphogroups/BM** |  |  |  |  |
| **Pairwise comparison** | **diff** | **lwr** | **upr** | **p adj** |
| 2P3MR-2P2MS | 0.25171249 | -0.33546 | 0.838885 | 0.757465 |
| 2P3MS-2P2MS | 0.10712009 | -0.40903 | 0.623274 | 0.978348 |
| 3P3ML-2P2MS | 0.97480036 | 0.402807 | 1.546793 | 6.66E-05 |
| 3P3MR-2P2MS | 0.89166805 | 0.366691 | 1.416646 | 7.11E-05 |
| 2P3MS-2P3MR | -0.1445924 | -0.65052 | 0.36134 | 0.932166 |
| 3P3ML-2P3MR | 0.72308787 | 0.160302 | 1.285874 | 0.004847 |
| 3P3MR-2P3MR | 0.63995557 | 0.125025 | 1.154886 | 0.007071 |
| 3P3ML-2P3MS | 0.86768027 | 0.379448 | 1.355913 | 2.91E-05 |
| 3P3MR-2P3MS | 0.78454797 | 0.352348 | 1.216748 | 1.87E-05 |
| 3P3MR-3P3ML | -0.08313231 | -0.58068 | 0.41442 | 0.990388 |
| **B: Morphogroups/P4** |  |  |  |  |
| **Pairwise comparison** | **diff** | **lwr** | **upr** | **p adj** |
| 2P3MR-2P2MS | 1.508548 | -0.01729 | 3.034386 | 0.054227 |
| 2P3MS-2P2MS | 0.45074 | -0.89055 | 1.792027 | 0.883723 |
| 3P3ML-2P2MS | 7.506366 | 6.019974 | 8.992757 | 0 |
| 3P3MR-2P2MS | 3.563341 | 2.199125 | 4.927558 | 0 |
| 2P3MS-2P3MR | -1.057808 | -2.37253 | 0.256916 | 0.175825 |
| 3P3ML-2P3MR | 5.997818 | 4.535351 | 7.460285 | 0 |
| 3P3MR-2P3MR | 2.054793 | 0.716684 | 3.392902 | 0.000415 |
| 3P3ML-2P3MS | 7.055626 | 5.786896 | 8.324356 | 0.0000000 |
| 3P3MR-2P3MS | 3.112601 | 1.989479 | 4.235723 | 0 |
| 3P3MR-3P3ML | -3.943025 | -5.23597 | -2.65008 | 0 |
| **C: Tooth number/BM** |  |  |  |  |
| **Pairwise comparison** | **diff** | **lwr** | **upr** | **p adj** |
| 4-5 | 0.1572848 | -0.25619 | 0.570758 | 0.639081 |
| 4-6 | 0.9245746 | 0.510042 | 1.339107 | 1.8E-06 |
| 5-6 | 0.7672898 | 0.475671 | 1.058909 | 0 |
| **D: Tooth number/P4** |  |  |  |  |
| **Pairwise comparison** | **diff** | **lwr** | **upr** | **p adj** |
| 4-5 | 0.8177346 | -0.58143 | 2.216902 | 0.350384 |
| 4-6 | 5.124122 | 3.721371 | 6.526873 | 0 |
| 5-6 | 4.3063873 | 3.319566 | 5.293208 | 0 |

