## Supplementary Methods S1 for "Bat teeth illuminate the diversification of mammalian tooth classes"

### Mathematical Modelling Supplementary Information for *Growth rate as a modulator of tooth patterning during adaptive radiations*

Alexa Sadier, Neal Anthwal, Andrew L. Krause, Renaud Dessales, Michael Lake, Laurent Bentolila, Robert Haase, Natalie Nieves, Sharlene Santana and Karen Sears

#### 1 Overview of Supplement

Here we propose a simple reaction-diffusion model of the formation and organisation of bat teeth primordia as the jaw grows. Between different species the size of the teeth and jaw, as well as the number and sequence of appearance of each bud, vary substantially. From its original appearance, these buds become the signalling centre of each tooth, and do not change as the jaw extends in each direction.

We will first consider sequences (in time) of insertions of each tooth bud, given by where it first appears in relation to the other teeth primordia. SI Figure 1 shows the experimentally-obtained sequences of insertions which come in four prototypical types across the observed species, though morphology, jaw length, and other details may change between each species even if the sequence of insertions is the same. The modelling presented here will be qualitative – as we do not know the details of the underlying molecular kinetics, we instead aim to provide examples where a simple reaction-diffusion theory can capture different sequences of tooth bud insertions over time, changing only aspects of the domain growth.

In Section 2 we present a simple reaction-diffusion model on a static domain to use as a basic framework for pattern formation, and discuss aspects of internal localisation of spots, representing the tooth buds. In Section 3 we extend this framework to include non-uniform domain growth, and propose a simple way of modelling such growth. We use this model in Section 4 to show how simple differential growth can capture a large variety of the insertion sequences observed, for equally sized teeth primordia. We then show in Subsection 4.3 how spatial heterogeneity (modelling other aspects of cell signalling) can in principle modulate size and spacing of teeth primordia, though at the cost of increasing model complexity (see [11] for discussions on the generality and limitations of such approaches). Finally in Section 5 some closing remarks about extending the basic ideas, particularly with an eye towards challenges in the theory, are made.

#### 2 Model in Static Domains (No Growth)

Here we present the basic model, using the classical Gierer-Meinhardt kinetics [2]. In the absence of detailed knowledge about the underlying signalling network, we consider a presumptive activator $u$ and inhibitor $v$ that obey the following system of reaction-diffusion equations,

$$\frac{\partial u}{\partial t} = \underbrace{\nabla^2 u}_{\text{Diffusion}} + \underbrace{\gamma + \frac{u^2}{v}}_{\text{Production}} - \underbrace{au}_{\text{Degradation}}, \tag{1}$$

$$\frac{\partial v}{\partial t} = \underbrace{d\nabla^2 v}_{\text{Diffusion}} + \underbrace{u^2}_{\text{Production}} - \underbrace{bv}_{\text{Degradation}}, \tag{2}$$

where $\gamma$ is a background or basal production of the activator, $a$ and $b$ are the degradation rates of the activator and inhibitor respectively, $d$ is proportional to the ratio of diffusivities, and $\nabla^2$ represents the Laplacian (diffusion) in a domain $\boldsymbol{x} \in \Omega \subseteq \mathbb{R}^n$, where we will consider two and three dimensional domains, so $n = 2, 3$. We remark that these equations are non-dimensonalized, so that these four parameters are really groupings of dimensional

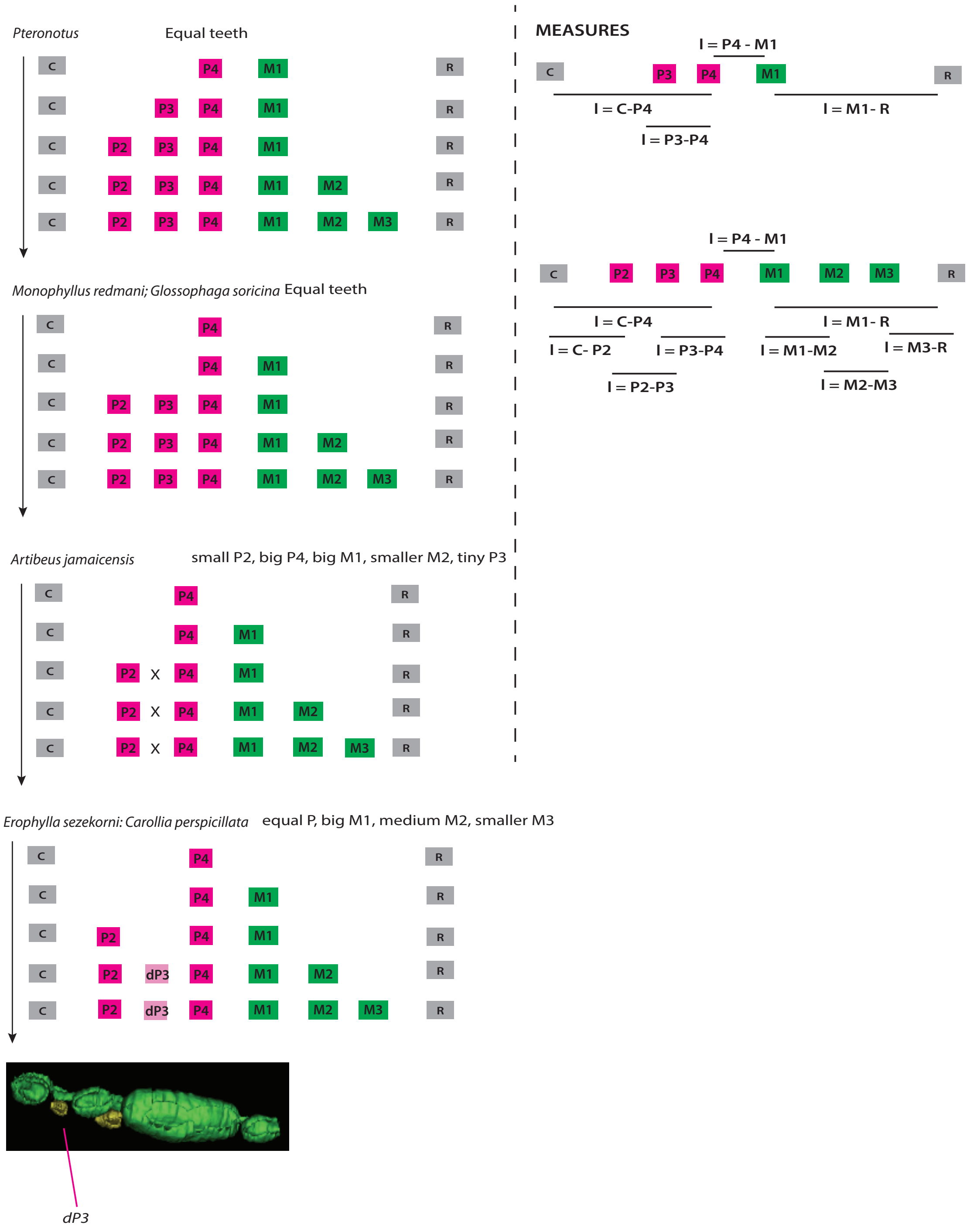


SI Figure 1: Sequence of primordia insertions as the jaw evolves across four different bat species, with time going downwards in each of the four panels.

parameters specific to the biochemistry involved. The non-dimensionalized form is taken such that the lengthscale of the domain $L$ is in units comparable to the activator's diffusion, which is taken to unity. Again, as we lack precise data on the presumptive morphogens, these may represent individual proteins, large signalling networks, or even cells in different stages of differentiation. Hence the model is not meant to capture quantitative details, but to test qualitative hypotheses of what underlies differences in morphology between species.

This system has a single spatially homogeneous steady state given by

$$u^* = \frac{b+\gamma}{a}, \quad v^* = \frac{(u^*)^2}{b} = \frac{(b+\gamma)^2}{a^2 b}.$$

We employed the usual Turing-type analysis to find parameters for which this steady state is stable in the absence of diffusion, but unstable to spatial perturbations when diffusion is considered, as these will typically lead to patterned solutions [8, 4]. In two and three spatial dimensions, within the parameter regimes identified above, the Gierer-Meinhardt system will generically form periodically arranged spots. The precise amplitude, wavelength, and arrangement of these spots depends on both the kinetic and diffusion parameters, as well as the geometry and boundary conditions. We primarily used mixed boundary conditions of the following form to prevent spots from accumulating near the boundaries:

$$u = 0, \quad \boldsymbol{n} \cdot \nabla v = 0, \quad \text{for all } \boldsymbol{x} \in \partial\Omega, \tag{3}$$

where $\boldsymbol{n}$ is the unit normal out of the domain, and $\partial\Omega$ represents the boundary. See [7] for a biological and mathematical discussion of these boundary conditions. The first of these conditions says that the activator concentration is fixed to zero at the edge of the region of interest (the jaw), and the inhibitor cannot diffuse outside of this region (i.e. a no-flux condition). For comparison in SI Figure 3(a), we also used the Neumann conditions,

$$\boldsymbol{n} \cdot u = 0, \quad \boldsymbol{n} \cdot \nabla v = 0, \quad \text{for all } \boldsymbol{x} \in \partial\Omega. \tag{4}$$

#### 2.1 Numerical Details

This model was simulated using commercial finite element software (Comsol Multiphysics V5.6) on rectangular and rectangular cuboid domains. Convergence checks in space and time were carried out, and specific simulations were also carried out in Matlab using the 5 (in 2D) and 7 (in 3D) point finite difference stencil (to ensure the finite element solutions were converging to the same solutions found via a different approach). In 2D, $50,786$ triangular finite elements were used, and in 3D $23,887$ tetrahedral elements were used, with solutions compared to those on finer meshes in both cases for particular simulations to ensure convergence. Initial data were taken as $u(\boldsymbol{x}, 0) = u^*\xi(\boldsymbol{x})$ and $u(\boldsymbol{x}, 0) = u^*\eta(\boldsymbol{x})$, where $\xi$ and $\eta$ are normally-distributed random variables (at each spatial grid point) with mean 1 and standard deviation $10^{-2}$. Larger and smaller random perturbations had little effect on the pattern sequence, as long as the timescales were sufficiently large. Even exactly homogeneous initial data form identical spatial patterns, as discussed in [7], due to the boundary conditions acting as a spatial heterogeneity. We therefore anticipate these results to be essentially insensitive of the initial conditions used.

#### 2.2 Simulation Results

In SI Figure 2 we plot two examples of static spot formation in two and three dimensional domains. The domain sizes here are fairly arbitrary, and lengthening or shortening them in the lateral direction would lead to more or fewer spots respectively. We also demonstrate the use of thresholding to indicate where plausible cell fate specification boundaries may be. This is particularly useful in the 3-D case for visualization purposes.

In SI Figure 3, we explore the use of Neumann (no-flux) boundary conditions (4) in panel (a), thin domains in (b), and thicker domains in (c). As shown in [7], no-flux boundaries can lead to spot formation along boundaries, where 'half-spots' may form, which is not something observed during bat teeth development but could be explained by other developmental and/or physico-chemical constraints present during the development of bat jaws. Such half-spot patterns motivate the use of the mixed boundary conditions (3). Thin domains using the mixed boundary conditions (3), shown in (b), exhibit elliptical spots rather than circular ones, though appear to be otherwise unchanged from simulations with larger lateral extents. If the lateral extent of the domain becomes too large,

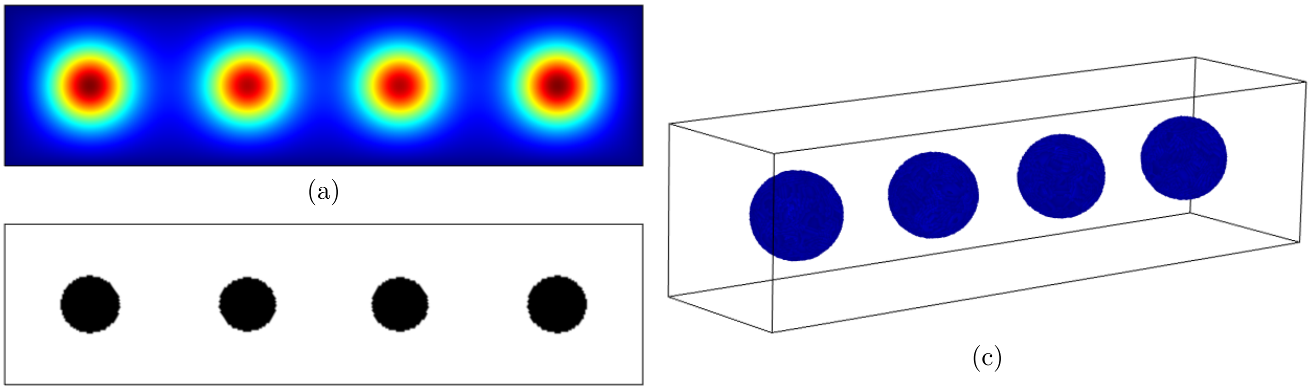

(a) (b) (c)

SI Figure 2: 2-D and 3-D simulations of (1)-(2) with $a = 0.5$, $\gamma = 1$, $b = 5.5$. The 2D plots are in a domain of size $15 \times 60$, and the 3D plot is in a domain of size $15 \times 15 \times 60$. (b) and (c) were 'thresholded' so that only the regions where $u > 25$ are shown, indicating a plausible cell fate specification boundary, though the actual concentration diffuses away from these peaks as shown in (a).

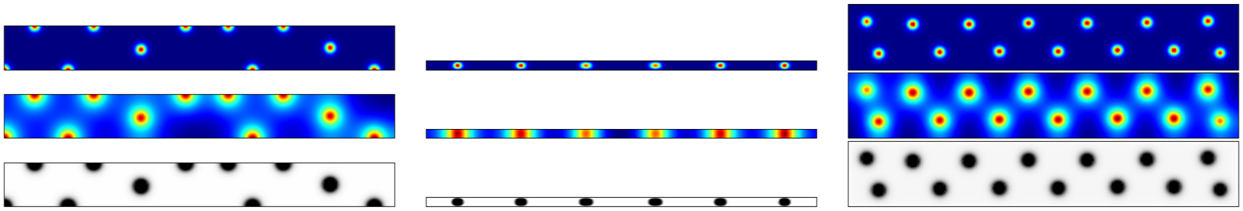

(a) Neumann BCs, $L_y = 2$ (b) Mixed BCs, $L_y = 0.4$ (c) Mixed BCs, $L_y = 3$

SI Figure 3: Simulations of the reaction-diffusion system with different geometries and boundary conditions. We show concentrations of the activator $u$ in the top row, inhibitor $v$ in the middle row, and a thresholded version of the activator in the bottom row where black is $u > u_{\text{Threshold}}$. The parameters used are $L_x = 18$, $L_y = 2$, $a = 0.5$, $b = 1$, $\gamma = 0.1$, $R = 0.3$, and $d = 0.005$. The initial length of the domain is $L_x = 4.5$, and the final length is $5L_x = 22.5$.

however, more than one row of spots may form, as in panel (c). This condition, which is not seen in mammals, is supposed to be due to a restriction of the mammalian dentition to one single row due to a pre-pattern, independently from reaction-diffusion mechanisms [12].

### 3 Non-Uniform Growth Model

Following [1], we formulate a model on a domain with non-uniform growth. We implement this by considering only growth in the $x$ direction, which we will rescale to an initial Lagrangian unit square (the rescaling in the $y$ direction is just a standard scaling, whereas in the $x$ direction the change of coordinates from a growing frame will introduce additional terms). As in [1], we consider a domain with two apically growing segments of possibly different growth rates, $r_1(t)$ and $r_2(t)$ (the first will represent the premolar region, and the second the molar region) according to the observations and mesearments made on $\mu$CT scans. We write $\xi$ as the rescaled $x$ coordinate and $y$ rescaled by length $L_y$ (noting that we think of $(\xi, y) \in [0,1]^2$). In the Eulerian frame then the total domain length is $L_x r(t)$ where $L_x$ is the initial length, and $r(t) = 1 + r_1(t) r_2(t)$.

We then have the evolution equations,

$$\frac{\partial u}{\partial t} = d\left(\frac{1}{(L_x r(t))^2}\frac{\partial^2 u}{\partial \xi^2} + \frac{1}{L_y^2}\frac{\partial^2 u}{\partial y^2}\right) + \left(\frac{\xi \dot{r}(t) - \dot{r_1}(t)}{r(t)}\right)\frac{\partial u}{\partial \xi} + \gamma + \frac{u^2}{v} - au, \tag{5}$$

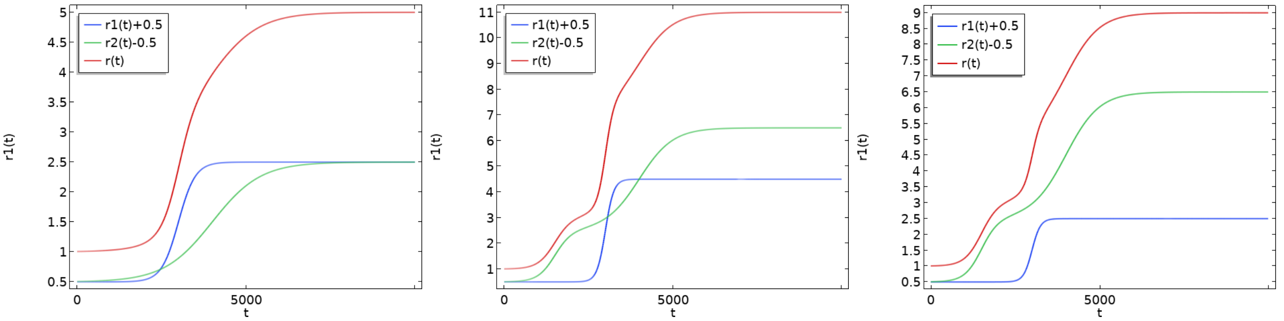


SI Figure 4: Growth functions in (a) for the domain given by $r_1(t) = 2B(t/T_f; 20, 0.3T_f)$, $r_2(t) = 1 + 2B(t/T_f; 7, 0.4T_f)$ in (b) for the domain given by $r_1(t) = 4B(t/T_f; 40, 0.3T_f)$, $1 + 4B(t/T_f; 10, 0.4T_f) + 2B(t/T, 20, 0.15T_f)$ and in (c) for the domain given by $r_1(t) = 2B(t/T_f; 40, 0.3T_f)$, where $T_f = 10^4$ is a final timescale, and $r(t) = r_1(t) + r_2(t)$. Note the shift by 0.5 in the plots is to center the domain at $r_1(0)$ and $r_2(0)$ so that $r_1(t)$ and $r_2(t)$ can be compared as $t$ increases.

$$\frac{\partial v}{\partial t} = \frac{1}{(L_x r(t))^2}\frac{\partial^2 v}{\partial \xi^2} + \frac{1}{L_y^2}\frac{\partial^2 v}{\partial y^2} + \left(\frac{\xi \dot{r}(t) - \dot{r_1}(t)}{r(t)}\right)\frac{\partial v}{\partial \xi} + u^2 - bv, \tag{6}$$

where we have the additional terms due to growth are given by the advection (first order space derivatives). For apical growth these essentially arise due to a change of coordinate system.

In principle this allows us to describe the non-uniform growth by specifying the two growth functions on each endpoint. More general nonuniform growth functions are difficult to determine, even if mitosis frequency can be measured across the jaw, and in general comparing patterns which form on timescales comparable with growth is quite difficult - much of their characteristics will depend on the nonlinearities chosen, which are impossible to know without a detailed understanding of the biochemical signalling. So we will build up sensible choices of growth functions to qualitatively match domain expansion, under this simple apical growth model.

We use the following smooth function with two parameters:

$$B(t; g, t_j) = \frac{1 + \tanh(g(t - t_j))}{2}, \tag{7}$$

where $g$ is a smoothing coefficient, and $t_j$ is a switching time. This function smoothly goes from 0 for small values of $t$, to 1 for larger values. Specifically, it is monotonically increasing and for $|g(t - t_j)| > 2.3$, $B$ is within 0.1% of 0 (for $t < t_j$) or 1 (for $t > t_j$). We can then vary $g$ to have the growth occur over longer periods of time, and take linear combinations of these $B$ functions to represent more complex sequences of growth.

### 4 Numerical Results Matching Sequence of Insertions

We now use the framework described to reconstruct the insertion sequences shown in SI Figure 1.

#### 4.1 Equally-Sized Teeth

We use simple combinations of the growth functions above to match the first three sequences of insertions in SI Figure 1. We aim only to match the sequence of the insertions, noting that quantitative aspects (such as matching a particular species' time in each stage) can be readily tuned.

We start by giving growth functions for *Pteronatus quadridens* in SI Figure 4(a), and show the sequence of insertions in SI Figure 5. As noted, the precise timings can be tuned by properly tuning the growth functions. We show sequences of insertions for *Monophyllus redmani* and *Glossophaga soricina* in SI Figure 6, using the growth functions given in SI Figure 4(b). Note that in SI Figure 6, we have included a snapshot of P3 forming near $t = 0.3T_f$, . The time frame where P3 exists but P2 does not can be made arbitrarily small with more rapid growth of $r_1(t)$ In SI Figure 7, we give the insertion sequence for *Artibeus jamaicensis*, in the absence of P3 and assuming that the primordia are equally sized and spaced.

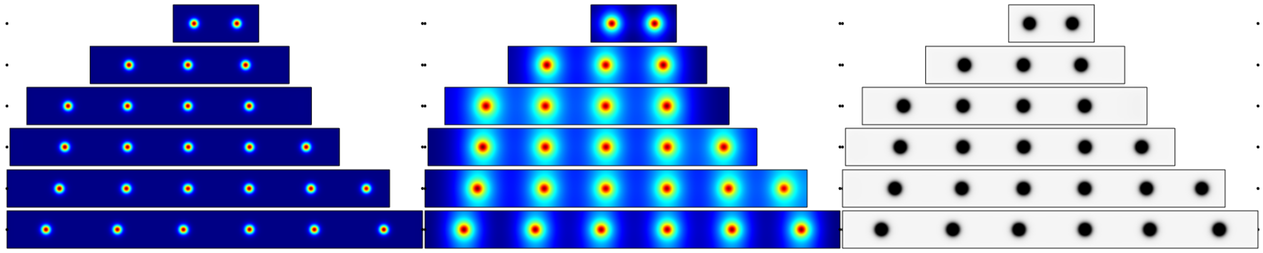

(a) $u$ (b) $v$ (c) $u > u_{\text{Threshold}} = 2$

SI Figure 5: Simulations of the reaction-diffusion system (5)-(6) with the evolution of the domain given in SI Figure 4(a). We show concentrations of the activator in (a), inhibitor in (b), and a thresholded version of the activator in (c). The top row is shown at time $T = 0.1T_f$, where $T_f = 10^4$ is a final timescale, and each subsequent row is shown at $T = 0.3T_f$, $T = 0.35T_f$, $T = 0.4T_f$, $T = 0.5T_f$, and $T = T_f$. The parameters used are $L_x = 4.5$, $L_y = 2$, $a = 0.5$, $b = 1$, $\gamma = 0.1$, $R = 0.3$, and $d = 0.005$. The initial length of the domain is $L_x = 4.5$, and the final length is $5L_x = 22.5$.

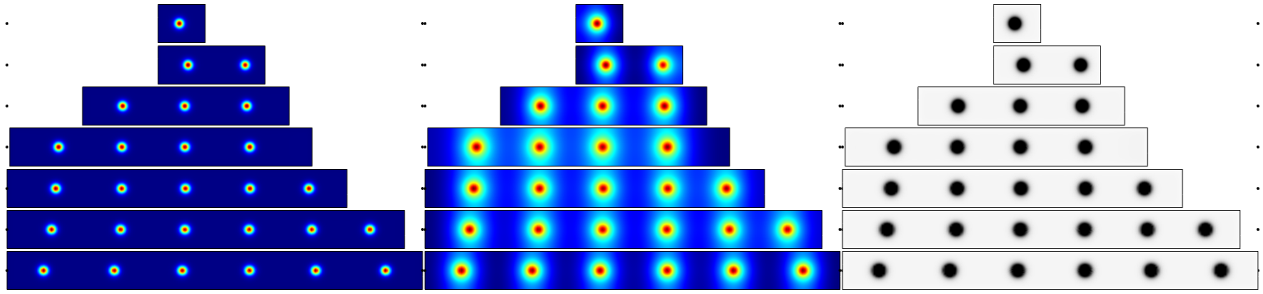

(a) $u$ (b) $v$ (c) $u > u_{\text{Threshold}} = 2$

SI Figure 6: Simulations of the reaction-diffusion system (5)-(6) with the evolution of the domain given in SI Figure 4(b). We show concentrations of the activator in (a), inhibitor in (b), and a thresholded version of the activator in (c). The top row is shown at time $T = 0.1T_f$, where $T_f = 10^4$ is a final timescale, and each subsequent row is shown at $T = 0.2T_f$, $T = 0.35T_f$, $T = 0.4T_f$, $T = 0.5T_f$, and $T = T_f$. The parameters used are $L_x = 2$, $L_y = 2$, $a = 0.5$, $b = 1$, $\gamma = 0.1$, $R = 0.3$, and $d = 0.005$. The initial length of the domain is $L_x = 2$, and the final length is $10L_x = 20$.

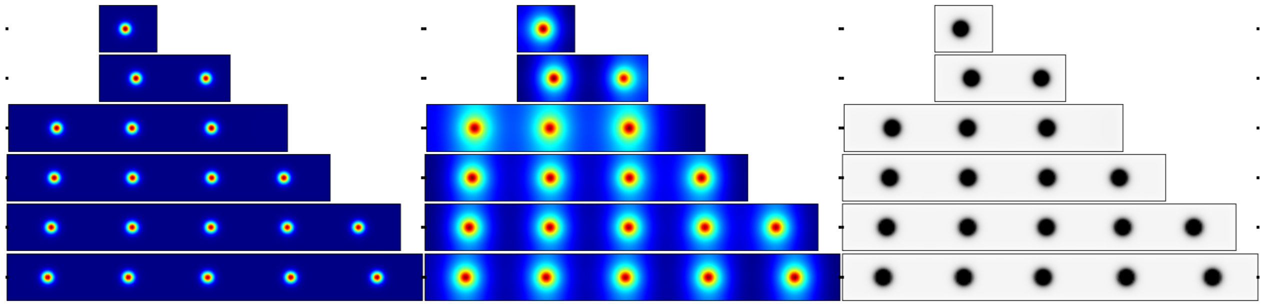

(a) $u$ (b) $v$ (c) $u > u_{\text{Threshold}} = 2$

SI Figure 7: Simulations of the reaction-diffusion system (5)-(6) with the evolution of the domain given in SI Figure 4(c). We show concentrations of the activator in (a), inhibitor in (b), and a thresholded version of the activator in (c). The top row is shown at time $T = 0.1T_f$, where $T_f = 10^4$ is a final timescale, and each subsequent row is shown at $T = 0.2T_f$, $T = 0.35T_f$, $T = 0.4T_f$, $T = 0.5T_f$, and $T = T_f$. The parameters used are $L_x = 2$, $L_y = 2$, $a = 0.5$, $b = 1$, $\gamma = 0.1$, $R = 0.3$, and $d = 0.005$. The initial length of the domain is $L_x = 2$, and the final length is $9L_x = 18$.

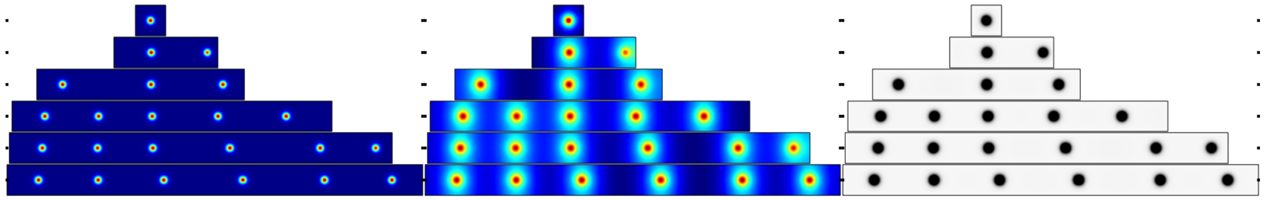


SI Figure 8: Simulations of the reaction-diffusion system (8)-(10) with the evolution of the domain given by $r_1(t) = 1 + 8.5B(t/T_f; 20, 0.3T_f)$ and $r_2(t) = 1 + 12B(t/T_f, 40, 0.38T_f) + 5B(t/T_f, 40, 0.25T_f)$. We show concentrations of the activator in (a), inhibitor in (b), and a thresholded version of the activator in (c). The top row is shown at time $T = 0.1T_f$, where $T_f = 10^4$ is a final timescale, and each subsequent row is shown at $T = 0.26T_f$, $T = 0.33T_f$, $T = 0.38T_f$, $T = 0.4T_f$, and $T = T_f$. The parameters used are $L_x = 2$, $L_y = 2$, $a = 0.5$, $b = 1$, $\gamma = 0.1$, $R = 0.3$, and $d = 0.005$. The initial length of the domain is $L_x = 2$, and the final length is $13.75L_x = 18$.

#### 4.2 Non-Apical Domain Growth Leads to Internal Insertions

In the fourth set of observed cases (*Erophylla sezekomi* and *Carollia perspicillata*), as well as the creation of new bud elements at the ends of the growing jaw, we also have the insertion of dP3 between two pre-existing buds. If we assume the domain growth is non-uniform, but also not purely apical, we can obtain a sequence of insertions matching this case. Again following [1], we consider a Lagrangian domain which is a unit square, and assume uniform growth at a rate $r_1(t)$ for $x \in [0, \theta)$, and uniform growth at a rate $r_2(t)$ for $x \in [\theta, 1]$. The total domain length in the $x$ direction at time $t$ is then given by $L_x r(t) = L_x(r_1(t)\theta + r_2(t)(1-\theta))$.

We can then write the equations of motion in the Lagrangian frame as,

$$\frac{\partial u}{\partial t} = d\left(\frac{1}{(L_x r(t))^2}\frac{\partial^2 u}{\partial \xi^2} + \frac{1}{L_y^2}\frac{\partial^2 u}{\partial y^2}\right) + \mathcal{B}(x)\frac{\partial u}{\partial \xi} + \gamma + \frac{u^2}{v} - au - 2S(x)u, \tag{8}$$

$$\frac{\partial v}{\partial t} = \frac{1}{(L_x r(t))^2}\frac{\partial^2 v}{\partial \xi^2} + \frac{1}{L_y^2}\frac{\partial^2 v}{\partial y^2} + \mathcal{B}(x)\frac{\partial v}{\partial \xi} + u^2 - bv - 2S(x)u, \tag{9}$$

where the functions $S$ and $\mathcal{B}$ are defined by,

$$S(x) = \begin{cases} \frac{\dot{r}_1(t)}{r_1(t)}, & \xi < \theta \\ \frac{\dot{r}_2(t)}{r_2(t)}, & \xi \geq \theta \end{cases}, \quad \mathcal{B}(x) = \begin{cases} \xi\left(\frac{\dot{r}(t)}{r(t)} - \frac{\dot{r}_1(t)}{r_1(t)}\right), & \xi < \theta \\ \xi\left(\frac{\dot{r}(t)}{r(t)} - \frac{\dot{r}_2(t)}{r_2(t)}\right) - \theta\left(\frac{\dot{r}_1(t)}{r_1(t)} - \frac{\dot{r}_2(t)}{r_2(t)}\right), & \xi \geq \theta \end{cases}. \tag{10}$$

We note that the $S$ term represents a dilution of morphogen due to the growth, which is neglected in the apical case as there the growth is so localized that no dilution occurs. The advection term, given by $\mathcal{B}$, is also more complicated in this case.

We plot the morphogen concentrations from our simulations in SI Figure 8. We remark that, besides the form of the growth and the final domain size, the parameters are identical to the other cases considered. Here the final domain is larger (so the buds appear smaller), as the dilution of concentration leads to overall fewer insertions on the timescale of growth. We observe the same sequence of insertions as given in the fourth case of SI Figure 1. Of course this sequence of insertions could be obtained in other ways by tuning parameters, or considering more complicated combinations of apical and locally-uniform regions of growth, but our aim here is to demonstrate that differential growth alone is sufficient to recapitulate these sequences.

#### 4.3 Heterogeneity Allows for Variation in Teeth Sizes

As shown above, all of the localized solutions (spots) observed in SI Figures 5-7 are approximately equally sized and spaced as the domain evolves. That is, they are all roughly circular and equally spaced, though the precise spacing can vary slightly as the domain expands (this can be understood as the domain is between two integer

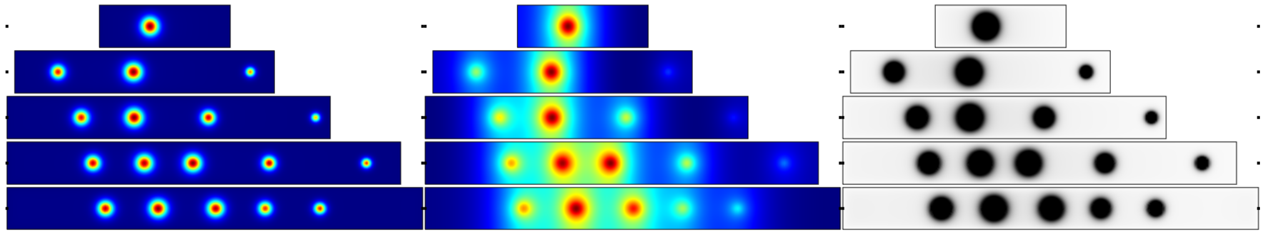

(a) $u$ (b) $v$ (c) $u > u_{\text{Threshold}} = 2$

SI Figure 9: Simulations of the reaction-diffusion system (12)-(13) with the evolution of the domain given in SI Figure 4(c). We show concentrations of the activator in (a), inhibitor in (b), and a thresholded version of the activator in (c). The top row is shown at time $T = 0.2T_f$, where $T_f = 10^4$ is a final timescale, and each subsequent row is shown at $T = 0.33T_f$, $T = 0.4T_f$, $T = 0.5T_f$, and $T = T_f$. The parameters used are $L_x = 1.8$, $L_y = 2$, $a = 0.5$, $b = 1$, $\gamma = 0.1$, $R = 0.3$, and $d = 0.005$. The initial length of the domain is $L_x = 1.8$, and the final length is $9L_x = 18$.

numbers of spots being permissible). Following studies of size and wavelength modulation in [9, 10] numerically and [6, 3] analytically, the authors in [11] demonstrated how simple spatial heterogeneities could be used to design relatively complex patterns. Here, we will introduce a simple modification of (5)-(6) in order to obtain sequences of primordia with different sizes.

We introduce the function,

$$A(\xi) = c_1 - c_2 \cos(\pi(\xi - c_3)), \tag{11}$$

which we use to modulate the linear degradation of the activator, $u$. This leads us to the equations,

$$\frac{\partial u}{\partial t} = d\left(\frac{1}{(L_x r(t))^2}\frac{\partial^2 u}{\partial \xi^2} + \frac{1}{L_y^2}\frac{\partial^2 u}{\partial y^2}\right) + \left(\frac{\xi \dot{r}(t) - \dot{r_1}(t)}{r(t)}\right)\frac{\partial u}{\partial \xi} + \gamma + \frac{u^2}{v} - aA(\xi)u, \tag{12}$$

$$\frac{\partial v}{\partial t} = \frac{1}{(L_x r(t))^2}\frac{\partial^2 v}{\partial \xi^2} + \frac{1}{L_y^2}\frac{\partial^2 v}{\partial y^2} + \left(\frac{\xi \dot{r}(t) - \dot{r_1}(t)}{r(t)}\right)\frac{\partial v}{\partial \xi} + u^2 - bv, \tag{13}$$

which are identical to (5)-(6) except for the term $aA(\xi)u$.

In SI Figure 9, we show a revised version of SI Figure 7 incorporating spatial heterogeneity in the activator kinetics as in (12)-(13), but using the same growth conditions as in SI Figure 5(c). We set $c_1 = 1.3$, $c_2 = 0.9$, and $c_3 = 0.4$. By design, P4 and M1 are larger, whereas P2 and M2 (again assuming P3 is absent) are smaller, and M3 is smaller still. We note that qualitatively similar patterns can be obtained with different values (or tweaking the 'base' kinetic parameters), particularly to exaggerate these size differences. However we note two barriers to this in general. Firstly, heterogeneity will typically modulate the wavelength between spots, and especially on growing domains this can lead to exaggerated spot movement (note that M1, M2, and M3 all emerge from the right boundary as very small spots before moving closer and becoming larger). Secondly, nonlinear effects can lead to complex spatiotemporal oscillations, and other bifurcations breaking the simple symmetries observed in the spatially homogeneous examples; see [5] for further discussion of this point.

### 5 Discussion

We have demonstrated how a very simple reaction-diffusion mechanism can produce most of the insertion sequences observed under a simple model of an apically growing domain. We have also shown that tuning kinetic parameters via spatial heterogeneity can match aspects of tooth modulation, at the cost of adding additional parameters and functions to the model, and influencing the wavelength (spacing) of teeth primordia. Of course, such modulation may actually occur via cell-fate specification downstream of a reaction-diffusion process which only fixes the signalling centres of each bud. Finally we also note that geometry can modulate these spot patterns, so that for thinner domains, elliptical patterns are obtained (see SI Figure 5(b)).
